# Understanding the role of IQ motif-containing GTPase-activating protein 1 as a potential disruptor of the essential adhesion module at cell-cell junctions

**DOI:** 10.64898/2026.10.22.761204

**Authors:** Nabanita Biswas, Amit Das

## Abstract

A key mechanism of stabilization of Cadherin-based cell-cell adhesions is through a mechanically modulated interaction to cortical actin filaments. This interaction is facilitated by a cytoplasmic adhesion module comprising of a complex of *β*-catenin and *α*-catenin. Recent evidence shows that under specific conditions the *β*-catenin/*α*-catenin complex gets disrupted because of mechanical disregulation of the junction maintenance machinery. The IQ motif-containing GTPase-activating protein 1 (IQGAP1) plays apparently a vital role as an internal regulator of the adhesion module, which competes for binding on *β*-catenin with *α*-catenin and is able to dissociate the latter from the adhesion complex. The underlying molecular mechanism is poorly understood with no clear evidence on what types of interactions are critical for a potential disruption. In this work, we combine steered-molecular-dynamics simulations and a multi-step protein–protein docking framework to dissect how mechanical detachment of *α*-catenin from the adhesion complex modulates the interactions between IQGAP1 and *β*-catenin. Our simulations show that as *α*-catenin detaches from *β*-catenin, IQGAP1 docks with progressively favorable scores. Interfacial analysis and subsequent mutagenesis studies, allow us to predict a molecular mechanism for an IQGAP1-driven disruption of the junctional adhesion module that centers on early onset of key favorable electrostatic interactions.

**Graphical Abstract:** Schematic representation of the force-dependent unbinding of *α*-catenin from the *β*-catenin/*α*-catenin complex, and step-wise docking of IQGAP1 to the deformed complex. Here the force of unbinding may come from high contractile forces on an actin filament attached to the complex. One possible source of the contractile force is strong and prolonged attachment to myosin motor proteins at high ATP concentrations.

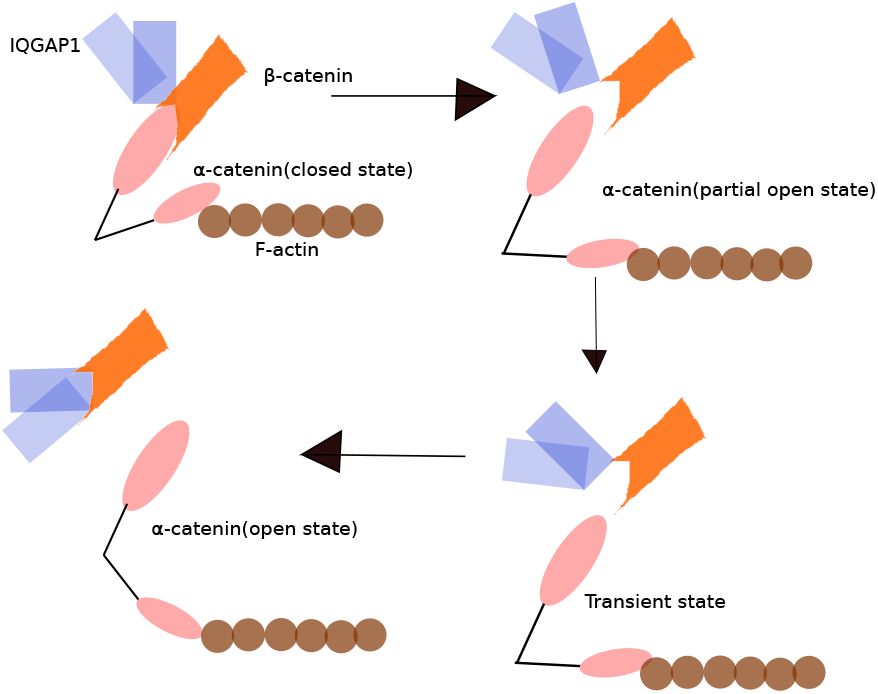

## 1. Introduction

Cell–cell adhesion is a fundamental biological process that enables cells within tissues to adhere to their neighbors through specialized surface molecules (Lodish et al. 2000). This ability to form stable yet dynamic contacts is essential for the development and maintenance of multicellular organisms (Takeichi 1991; Shawky and Davidson 2015). Adhesive interactions do more than simply connect adjacent cells; they provide a structural and signaling framework that supports tissue organization, integrity, and homeostasis (Gumbiner 1996; Nelson, Dickinson, and Weis 2013). In the absence of such adhesive mechanisms, complex tissue architecture would disintegrate, impairing key biological processes including embryonic development (Gilbert 2000), immune cell trafficking (Harjunpää et al. 2019), and wound repair (Peña and Martin 2024). Thus, cell-cell adhesion represents a foundational principle of multicellular biology, linking structural cohesion with dynamic regulation of cell behavior.

At the molecular level, cell–cell adhesion is mediated by several distinct families of transmembrane adhesion molecules, including cadherins, selectins, integrins, and IgSF-CAMs (Gumbiner 1996). Among these, cadherins are particularly central to tissue architecture, with E-cadherin-based adherens junctions serving as a model for understanding adhesive strength and mechanotransduction (Takeichi 1991). E-cadherin, a calcium-dependent cadherin protein, is essential for maintaining epithelial structure (Berx and Roy 2001; Hartsock and Nelson 2008). Its loss or dysfunction at cell–cell junctions leads to developmental abnormalities (Edelman 1983) and is strongly linked to cancer progression and epithelial-to-mesenchymal transition (EMT) (Nieto et al. 2016; Youssef and Nieto 2024).

E-cadherin mediated cell–cell adhesion relies on a highly organized supra-molecular architecture, assembled through a strongly regulated dynamic process (Bachir et al. 2017; Takeichi 2014). The E-cadherin extracellular domain forms *trans*–homophilic interactions with E-cadherin molecules on neighboring cells, while the cytoplasmic domain of E-cadherin interacts with *β*-catenin and *β*-catenin, in turn, interacts with *α*-catenin that link the cellular adhesion complex (E-cadherin cytoplasmic domain–*β*-catenin–*α*-catenin) to the actin cytoskeleton (Yamada et al. 2005). This E-cadherin–catenin–actin complex not only provides strong adhesive strength but also converts adherens junctions into hubs for signalling and cytoskeletal regulation (Nelson and Nusse 2004). Disruption of this cellular adhesion complex is frequently associated with pathology: cancer cells often lose E-cadherin expression during EMT, enabling increased motility and invasiveness (Onder et al. 2008; Thiery 2002; Jin et al. 2015; Mataraza et al. 2003; Yerramilli et al. 2022). A stable interaction between E-cadherin, *β*-catenin and *α*-catenin, particularly within the *β*-catenin N-terminal region (residues 120–151) is a critical element of junctional integrity. Perturbation of this *β*-catenin and *α*-catenin interface weakens tissue cohesion.

A wide range of biochemical and cellular evidence demonstrates that IQ motif-containing GTPase-activating protein 1 (IQGAP1) is a potent regulator of the *β*-catenin/*α*-catenin interaction and competes directly with *α*-catenin for binding to similar region on the N-terminal of *β*-catenin (Kuroda et al. 1998; Fukata et al. 1999). Fukata and co-workers have performed a comprehensive study in this regard (Fukata et al. 1999). They investigated whether IQGAP1 induces the dissociation of *α*-catenin from the *α*-catenin/*β*-catenin complex using an *in vitro* assay. They immobilized full length *β*-catenin and the *α*-catenin binding domain of *β*-catenin separately on affinity beads. These beads were then incubated with α-catenin and then eluted with IQGAP1 and two different control proteins. They confirmed *α*-catenin dissociation through its detection in the eluates by SDS-PAGE only in case of IQGAP1, not the control proteins. They interpreted this to be a strong indication that IQGAP1 and *α*-catenin compete to bind *β*-catenin. Later they confirmed the competitive binding through separate bead-based assays with both IQGAP1 and α-catenin present when they show signatures of competition. Finally, their cell dissociation assays further confirm that IQGAP1 expression indeed weakens cell–cell adhesion by increasing sensitivity to mechanical disruption. Similar findings are reported in earlier studies (Bashour et al. 1997; Kuroda et al. 1998) and later confirmed by Fukata and co-workers (Fukata et al. 1999). Structural mapping shows that both IQGAP1 and *α*-catenin bind overlapping regions within *β*-catenin residues 120–151 (Aberle et al. 1994), providing a mechanistic basis for competitive displacement. What makes this competition so relevant is the observation that IQGAP1 is overexpressed in multiple cancers and promotes *β*-catenin stability and tumor progression (Jin et al. 2015; Mataraza et al. 2003; Mohapatra and Dixit 2022).

Collectively, these observations highlight IQGAP1 as a significant modulator of adherens junction stability with consequences for mechanotransduction (Noritake et al. 2005; Nouri, Timson, and Ahmadian 2020). The *β*-catenin/*α*-catenin complex forms the primary mechanical linkage between E-cadherin and actin, enabling cells to sense and respond to tension (Yap, Duszyc, and Viasnoff 2018). From above literature study one may argue that, as IQGAP1 weakens the *β*-catenin/*α*-catenin complex by competing for an overlapping interface on *β*-catenin with *α*-catenin, then a much smaller mechanical force would be sufficient to detach *α*-catenin from *β*-catenin. Conversely, if a junction is already weakened due to accumulation of myosin contractile load (Duque et al. 2024), it may be subjected to destabilization by IQGAP1. Literature on junctional mechanics suggests that *α*-catenin undergoes force-dependent conformational changes that modulate the stability of the adhesion module (Buckley et al. 2014; Windgasse and Grashoff 2025). Increased IQGAP1 expression has been associated with enhanced junction turnover and reduced epithelial cohesion (Noritake et al. 2005), indirectly supporting the idea that IQGAP1 may accelerate mechanical unbinding of the *β*-catenin/*α*-catenin complex at adherens junctions.

Some gap still remains regarding the molecular-level understanding of IQGAP1-driven dissociation of *α*-catenin from the complex with *β*-catenin. Previous experimental evidence involving various fragments of IQGAP1 have shown that the entire IQGAP1 structure may be essential for adequate binding to *β*-catenin and dissociation of *α*-catenin (Kuroda et al. 1998). Those data, along with later observations (Elliott, Allen, and Timson 2012), indicate that specific fragments of IQGAP1, especially the carboxy terminal, still exhibit dose dependent competiive binding to *β*-catenin. Majority of subsequent studies have focused on the overall regulation of IQGAP1 activity (Owen et al. 2008; Nouri, Timson, and Ahmadian 2020), not specifically IQGAP1/*β*-catenin interactions. Therefore, any further details of a key hotspot on IQGAP1 that could help us understand the molecular mechanism behind the above-mentioned competitive binding of IQGAP1 to *β*-catenin has been lacking. The sheer size of the IQGAP1 molecule (*∼*1700 residues) adds further complications to the problem.

In this work, we address this problem by considering the premise of ‘mechanically weakening’ of the *β*-catenin/*α*-catenin complex. Instead of a direct competitive binding model, we focus on an alternative, computational framework centered on a stage-wise displacement and multistep docking. Our approach integrates various computational techniques with atomistic details like, regular molecular dynamics (MD), steered molecular dynamics (SMD) to simulate different stages of the forced-displacement of *α*-catenin from the adhesion complex. Then we dock IQGAP1 to various stages of the deformed complex in steps, starting from partially attached to completely detached *α*-catenin. Our results indicate that binding of IQGAP1 to the deformed *β*-catenin/*α*-catenin complex becomes progressively favorable. Comparative interfacial analysis on three different docking protocols (fully blind, partially blind and restrained) help us predict specific amino acid pairings between IQGAP1 and *β*-catenin that could get established at very early stages of deformation of the above complex and lead to favorable binding. These two sets of evidence allow us to predict some residues on IQGAP1 as possible candidates for binding of *β*-catenin, which could be relevant for a potential IQGAP1-induced disruption of the adhesion complex. Our findings offer a quantitative framework to understand the molecular basis of how IQGAP1 perturbs cell-cell adhesion complex stability.

## 2. Materials and Methods

Our systematic workflow is described in Fig. 1. First, the initial configuration of the *β*-catenin/*α*-catenin complex are prepared using fragments of *β*-catenin and *α*-catenin from an available structure of the complex in the Protein Data Bank (PDB). These fragments (Fig. 2a-c, represent the interfacial region of the two proteins when they interact (Fig. 2c). Similarly, an available PDB structure of IQGAP1 fragment is chosen (Fig. 3a-b). Next, we deform the *β*-catenin/*α*-catenin structure by pulling on the *α*-catenin fragment using a SMD protocol (Lemkul and Bevan 2010). Then stepwise docking of IQGAP1 fragment is performed at different deformed configurations of the complex sampled at different time points of the pulling simulation. Each of these docking exercises are guided by a fixed set of active residues obtained from an independently docked *β*-catenin/IQGAP1 complex shown in Fig. 3b. Below, we describe the entire workflow in detail.

**Figure 1.:**
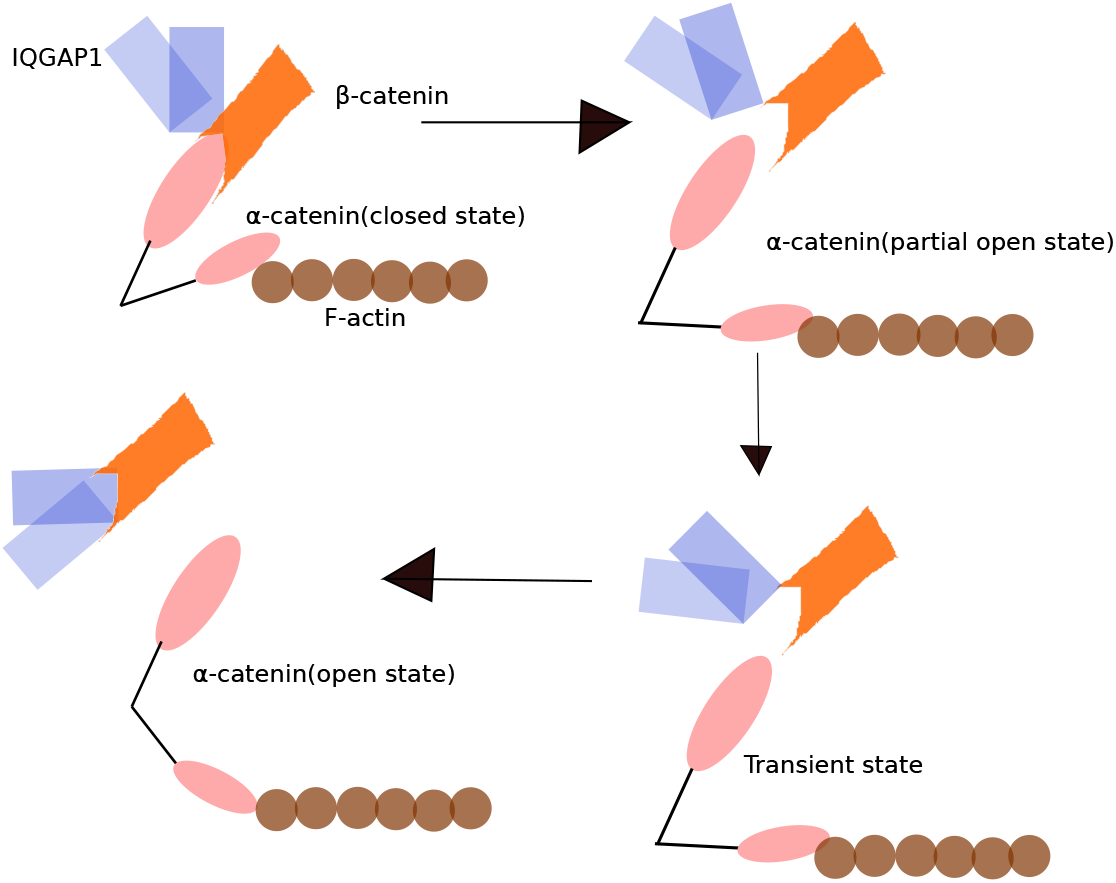
Schematic representation our workflow based on SMD: Steps of force-dependent unbinding of *α*-catenin from the *β*-catenin/*α*-catenin complex are shown. Different steps are distinguished by distinct conformational states of *α*-catenin. We dock IQGAP1 on several pulled conformers as shown here. Here the *α*-catenin is shown to be pulled by an actin filament under heavy contractile load from myosin motors.

**Figure 2.:**
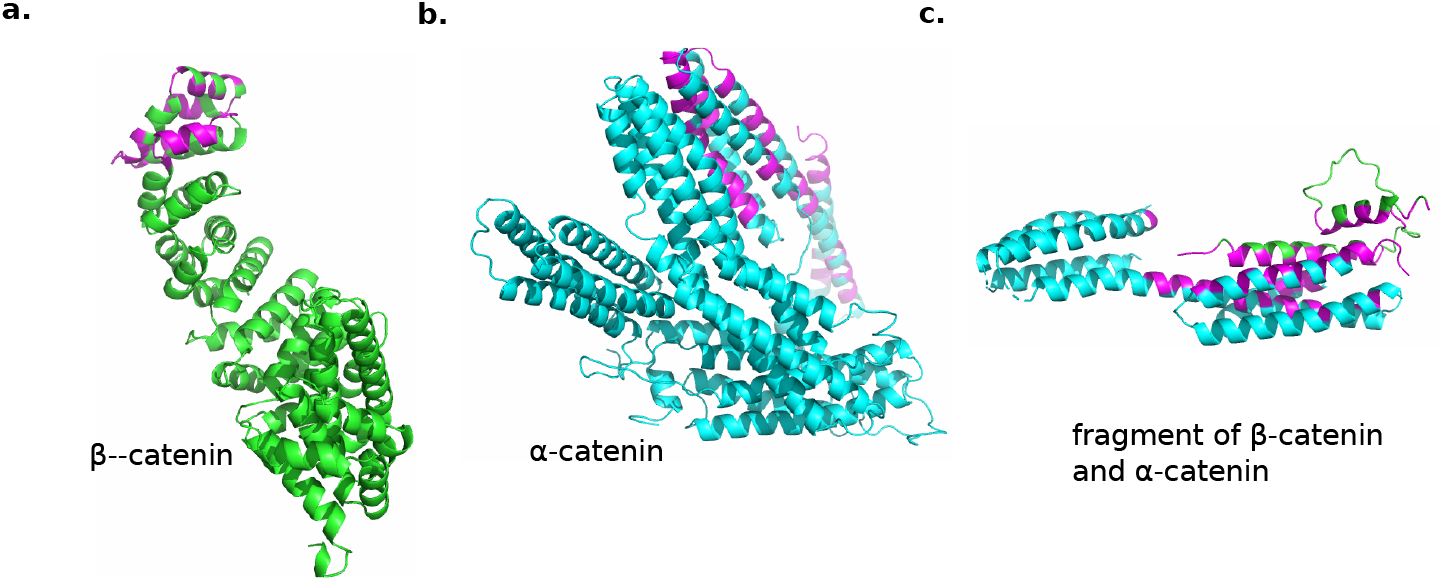
The structures of binding fragments of *β*-catenin and *α*-catenin that form the core of the adhesion complex. Complete structures of (a) *β*-catenin with the fragment highlighted, (b) *α*-catenin with the fragment highlighted, and (c) just the binding fragments with interfacial residues highlighted. The structures shown in a and b are from AlphaFold database (Fleming et al. 2025) and PDB ID 4IGG, respectively. The structure in panel c is from PDB ID 4ONS.

**Figure 3.:**
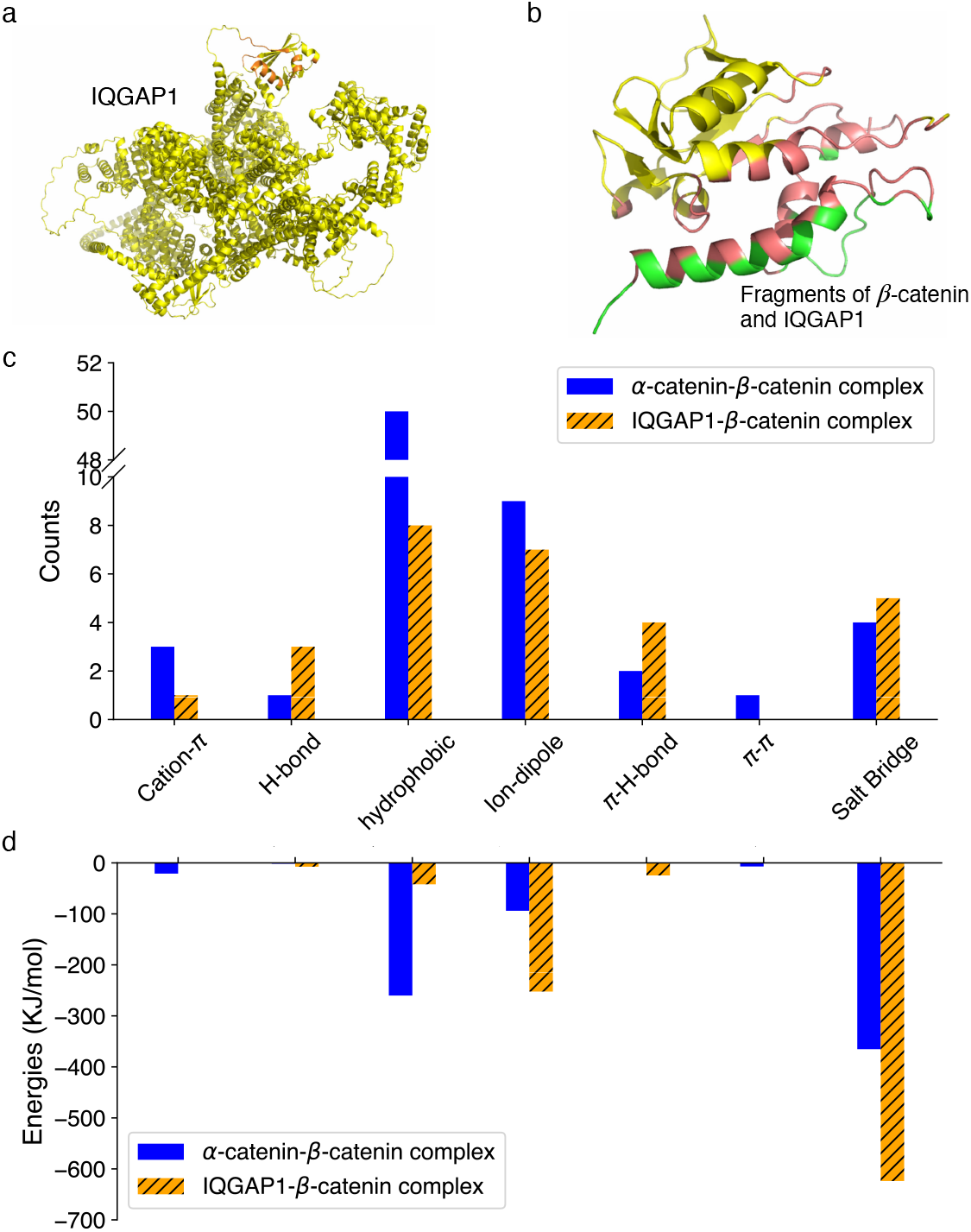
Comparison of the binding interfaces on *β*-catenin for *α*-catenin and IQGAP1. (a) Complete structure of IQGAP1 obtained from AlphaFold database (Fleming et al. 2025) with the binding fragment from PDB ID 1X0H highlighted. This is provided only as a guide to show the location of the fragment used for studies in this work. (b) The *β*-catenin/IQGAP1 binding fragments with interfacial residues highlighted. (c-d) The comparison of *β*-catenin/*α*-catenin interface and the *β*-catenin/IQGAP1 interface in terms of contributions from different types of inter-amino acid interactions. The numbers of various interaction types detected with a 4 Å distance cutoff on the side-chain-side-chain separation are shown in panel c. The corresponding energies are compared in panel d.

### 2.1. Protein Structure Preparation

The starting coordinates of the binding protein fragments in the *β*-catenin/*α*-catenin complex are chosen from the PDB ID 4ONS. The species of *α*-catenin in this reported structure is *α*-N-catenin, which is different from *α*-E-catenin that is usually associated with E-cadherin based cell-cell junctions. However, this structure is eventually used because, this is the only experimentally determined structure of *β*-catenin/*α*-catenin complex that we could find. A comparative study of the *α*-N-catenin sequence vs *α*-E-catenin sequence reveals *∼* 82% sequence identity that is also reported earlier (Ishiyama et al. 2013). The relevant region that interacts with *β*-catenin is the N-domain, which between the two varieties of *α*-catenin maintains a even higher homology according to previous reports (Ishiyama et al. 2013; Huber, Krohn, and Kemler 1997). Subsequently, we use BLASTP to compare a 27-amino acid segment of *α*-E-catenin (residues 117-143) that is necessary and sufficient for binding to *β*-catenin, as reported earlier (Huber, Krohn, and Kemler 1997). PDB ID 6O3E (same organism, Mus Musculus) is used for the reference sequence of *α*-E-catenin. This exercise shows that there is only a single residue difference (ASN to THR at position 123) when compared with the same region on *α*-N-catenin from 4ONS. This gives us confidence that our findings would remain valid even in case of E-cadherin-based epithelial adherens junctions involving *α*-E-catenin. We use the structure of IQGAP1 fragment from PDB ID 1X0H.

Necessary H-atoms are added to the protein structures to implement the physiological pH condition (pH = 7.4) (Dolinsky et al. 2004), ensuring correct protonation states of ionizable residues. The AlphaFold database for protein structures (Fleming et al. 2025) is used to obtain the complete structures of *β*-catenin, *α*-catenin and IQGAP1. This exercise is purely just to visualize the locations of the binding fragments on the respective protein-bodies. None of the AlphaFold-predicted structures are used for our subsequent calculations. To compute all the interatomic potential energies and forces in our molecular dynamics (MD) simulations (regular and steered), we choose the widely-used OPLS-AA force field (Jorgensen, Maxwell, and Tirado-Rives 1996), which offers a good description of both backbone and side-chain conformations of the amino acids. Subsequently, we solvate the protein fragments in a cubic box full of water molecules represented via the SPC model (Berendsen et al. 1981). A periodic buffer of 10 nm is applied in all of X, Y and Z directions to ensure that the proteins do not interact with their periodic images during the simulations, even in the pulled state. Adequate numbers of Na^+^ and Cl*^−^* are then introduced in the simulation box as counter-ions to make the system charge-neutral. The effective concentration of both the ions used in the simulation box turns out to be approximately 99 mM, which corresponds to an ionic strength of 99 mM (Solomon 2001). This value is close but on the lower-side compared to recently measured cellular ionic strength in range 110-130 mM (Liu, Poolman, and Boersma 2017).

### 2.2. Molecular Dynamics

We perform MD simulations to investigate the conformational stability of the different protein complexes involved in the study using GROMACS 2023.2 (Berendsen, van der Spoel, and van Drunen 1995; Abraham et al. 2015; Páll et al. 2020). Prior to the MD, potential energy of the system is minimized using the steepest-descent energy minimization process to remove steric clashes, unfavorable contacts, and other geometric inconsistencies introduced during protein structure preparation. Following minimization, the system is equilibrated in the constant number of particles (N), pressure (P), and temperature (T) or NPT ensemble under periodic boundary conditions to bring the temperature at 310 K and pressure at 1 Bar for a short time period of 100 ps using the Parrinello–Rahman barostat (Parrinello and Rahman 1981). During equilibration, positional restraints are employed to the heavy atoms of the protein backbone to allow the solvent molecules and ions to relax around the biomolecule without disrupting the protein’s initial conformation. After equilibration, we perform unrestrained production MD simulations on the protein complexes. For the *β*-catenin/*α*-catenin fragment complex our run length is 100 ns. Any loong-range effects from the electrostatic interactions are handled via the particle mesh Ewald (PME) method (Essmann et al. 1995) with a real-space cut-off of 1.0 nm, while van der Waals interactions are truncated at the same distance. All H-bonds are constrained using the LINCS algorithm (Hess et al. 1997), allowing stable integration of the equations of motion at the chosen time step of 2 fs. We also apply long-range dispersion corrections to energy and pressure to account for interactions beyond the cut-off. Throughout the simulation, various state parameters like temperature, pressure, total energy, and root-mean-square deviation (RMSD) of the protein backbone are monitored. Finally, we implement Steered-MD on the *β*-catenin/*α*-catenin fragment complex, to simulate detachment of *α*-catenin, on the randomly sampled snapshots chosen from the terminal part of the 100 ns long regular MD trajectory. For the wild type and mutant IQGAP1/*β*-catenin fragment complexes the MD run length is set to 200 ns. In each case we generate four replicates.

### 2.3. Steered Molecular Dynamics

The main backbone of our workflow is a force-dependent unbinding of *α*-catenin from the *β*-catenin/*α*-catenin complex. To realize this, we use a well known SMD protocol implemented via the non-equilibrium pulling method within GROMACS (Lemkul and Bevan 2010). In our setup, we employ a constant-velocity pulling on *α*-catenin as follows. *β*-catenin was position-restrained to mimic its stable membrane-anchored connection via E-cadherin. Conversely, *α*-catenin was tethered to the pulling spring to simulate the tensile load typically exerted by the actin cytoskeleton. The force constant of the virtual spring is set at 1000 kJ mol*^−^*^1^ nm*^−^*^2^, to ensure sufficient stiffness to capture unfolding events, while avoiding any artifacts from the restraint forces. The pulling direction is fixed to be along the principal molecular axis joining the centers of mass of the two protein fragments. We pull at two different velocities: 10 nm ns*^−^*^1^ and 12 nm ns*^−^*^1^. The current range of chosen velocities represents a reasonable trade-off between computational feasibility and non-equilibrium effects for SMD simulations. Essentially, such high pulling velocities are chosen only to simulate the mechanical weakening or complete detachment of the *β*-catenin/*α*-catenin complex in cost efficient manner. For both pull velocities, the SMD trajectory is sampled with a uniform periodicity of 2.3 ps to extract 500 conformers from a run length of 1.15 ns following the 100 ns regular MD run for subsequent analyses. Four replicates of each SMD trajectory are generated at both velocities by changing the initial random number seed. All replicates showed same qualitative trends of detachment dynamics of *α*-catenin.

From the changes in intermolecular contacts at the interface, we compute the time evolution of the *β*-catenin/*α*-catenin shared interface area during the pulling process using the PyMOL package (Schrödinger 2015). The interface area is defined as the set of atomic contacts between two molecules (chains or subunits) that leads to a loss of solvent-accessible surface area for both partners when they form a complex. We represent the interface area as half of the total decrease in accessible surface area when the two isolated components are brought together, using the formula: interface area = (*T_A_* + *T_B_ − T_AB_*)*/*2, where *T_A_* and *T_B_* are the solvent-accessible surface areas of the separate molecules and *T_AB_* is that of the complex.

We used the open-source visual molecular dynamics (VMD) package (Humphrey, Dalke, and Schulten 1996) version 2.0.0 (Gomes et al. 2026) for all visualization and structural inspection of the conformational states during the pulling and for generating images afterwards. Figures of proteins and residue-level interactions are generated using the PyMOL package.

### 2.4. Protein–Protein Docking

Apart from the SMD, we implement serial protein–protein docking to predict structural insights into the binding of IQGAP1 to the deformed *β*-catenin/*α*-catenin. Our docking experiments are performed using the High Ambiguity Driven protein–protein Docking (HAD-DOCK) version 3.0 (Van Zundert et al. 2016; Dominguez, Boelens, and Bonvin 2003; Giulini et al. 2025). HADDOCK uses various interaction restraints, derived from known or predicted binding residues, to guide the docking search. The latest modular version, also called HAD-DOCK3 (Giulini et al. 2025), allows flexibility of various independent refinement modules, namely an energy minimization, interface optimization and an integrated MD-based refinement in explicit solvent.

Our multi-step docking protocol is organized as the following: First, we identify different stages of the detachment of *α*-catenin fragment from the *β*-catenin fragment using SMD. From the 1.15 ns SMD trajectory, we have chosen a series of partially detached conformations of the *β*-catenin/*α*-catenin complex. These are referred to as the ‘conformers’. Our sampling periodicity is 2.3 ps. Accordingly, we choose total of 500 conformations. Thus, the conformation chosen at 23 ps is named as ‘conformer 10’, that at 230 ps as ‘conformer 100’, at 575 ps as ‘conformer 250’ and so on. Subsequently, we choose 15 conformers for docking.

For each conformer we dock the IQGAP1 fragment on the partially intact complex. Three different versions of the multi-step docking protocol are executed: a completely blind protocol, a partially blind protocol and a restrained protocol. For the completely blind protocol, the IQGAP1 fragment is permitted to bind anywhere on the *β*-catenin/*α*-catenin complex without any spatial or ambiguous interaction restraints (AIRs). In the partially blind case, the IQGAP1 fragment is permitted to bind anywhere on the *β*-catenin fragment. This is achieved by providing all possible residue combinations on the *β*-catenin fragment and the IQGAP1 fragment as the AIRs. In the restrained protocol, the IQGAP1 fragment is guided by the chosen active residues on *β*-catenin, included as AIRs. In absence of any experimental data on the specific interacting residues between *β*-catenin and IQGAP1, to the best of our knowledge, the active residues are obtained from an independently docked IQGAP1/*β*-catenin complex, highlighted in Fig 3. The interfacial data from this docking are listed in supplementary Table S2.

Each of our docking simulations provide a HADDOCK score for each model, which is an weighted sum of various energy terms, like electrostatic energy, van der Waals intermolecular energy, desolvation energy, various restraint energies, etc. and buried surface area. These scores essentially are used to rank various docked poses obtained for a given complex. HAD-DOCK3 also computes a DockQ value (Basu and Wallner 2016) and an interface RMSD and for each docking model, which represents a quality measure for protein-protein docking simulations. Typically such values are generated with respect to a reference structure provided in the calculation. In the current work, due to unavailability of any experimentally determined structure of the IQGAP1/*β*-catenin reference complex, HADDOCK3 considers the highest scoring model as a pseudo reference structure (Giulini et al. 2025) in each docking simulation. We monitor DockQ and interface RMSD values to assess the extent of convergence of the docking models relative to the internally selected reference model. Finally, we choose the top ten models for subsequent interfacial analyses.

The docking protocol is executed on the conformers from all four replicates for a given SMD trajectory. We inspected the interfacial interactions and found many of them consistently forming in majority of the replicates within reasonable variation. Therefore, the reported docking scores are calculated for each conformer by averaging the scores of the best models from four different replicates.

### 2.5. Interface and Mutational Analysis

To identify the molecular determinants of IQGAP1/*β*-catenin interactions, we analyze the interfaces of the top-ranked docking clusters. We adjudge any residue located within 4 Å of the partner protein as interface residues, as this cut-off captures both direct hydrogen bonds and van der Waals contacts. This is done using a combination of custom PyMOL routines (Vertrees 2015) and in-house python scripts. The interface residues are categorized in terms of their physico-chemical properties and potential interaction partners into several groups, namely ionic – forming salt bridges through electrostatic interactions with an oppositely charged residue, polar uncharged – that engage in H-bonds, residues with aromatic rings – *π* electron-donors to cations and dipoles, and non-polar residues – that primarily form hydrophobic pockets. The inter-amino acid interaction energies for the interfaces of the *β*-catenin/*α*-catenin and IQGAP1/*β*-catenin complexes are also estimated using the INTAA webserver (Vymětal et al. 2021) (see more details in section 1, supplementary text).

To probe the role of the any identified candidate residues in complex formation with moderate to high occurrence frequencies across the partially blind and restrained protocols, we introduce targeted point mutations in IQGAP1. Several charged residues are mutated to alanines, thus removing the associated salt-bridges and ion-dipole interactions. *In silico* mutagenesis is performed directly on the docked complexes using the MODELLER program (Šali and Blundell 1993; Webb and Sali 2016), followed by energy minimization and a short MD simulation to relax local geometries and optimize the conformation of the mutated side-chain. The docking scores for all the complexes of *β*-catenin/IQGAP1 mutants are estimated using HADDOCK3 and monitored as the differential score: Δ Score = (wild type score - mutant score). Finally, 200 ns long MD simulations are performed on the *β*-catenin/mutated IQGAP1 complexes, and their RMSD are compared with that of *β*-catenin/wild type IQGAP1 complex in order to get insight on the role of the key residues in stability of the complex.

## 3. Results

### 3.1. Competitive Binding Interface of α-catenin and IQGAP1 on β-catenin

To understand how strongly IQGAP1 might bind to *β*-catenin, we compare the binding interface of two protein fragment complexes: the *β*-catenin and *α*-catenin complex (Fig. 2c) and the *β*-catenin and IQGAP1 complex (Fig. 3b). We use the INTAA webserver (Vymětal et al. 2021) to analyze the two interfaces as described in supplementary text. The server outputs are analyzed as the following: the interfacial residues are identified by side-chain-side-chain contacts with a 4 Å distance cutoff. The residue list for the two complexes are included in Supplementary Tables S1 and S2. The energies for each identified interaction are also recorded and mentioned in the tables. We estimate an interaction energy of *≈ −*10 kJ mol*^−^*^1^ per residue for the *β*-catenin/*α*-catenin interface, whereas the IQGAP1/*β*-catenin interface shows *≈ −*32 kJ mol*^−^*^1^ per residue. This difference is striking given the former interface have nearly 70 contacts, with latter showing only 29. Such high stabilization for the salt bridges observed for IQGAP1-fragment could be indicative of these interactions to be protected from solvent exposure or external perturbations.

We also inspect the specific inter-amino acid interaction types at the two interfaces (Fig. 3c-d). Our analysis reveals that *α*-catenin and IQGAP1 fragments share an overlapping binding site on the *β*-catenin fragment and exhibit certain similar type of interactions. Both interfaces are supported by a number of hydrophobic and electrostatic contacts. The *β*-catenin/*α*-catenin complex features a significant number of hydrophobic interactions, about 50 accordingly to our analyses (supplementary table S1), whereas the *β*-catenin/IQGAP1 interface shows a fraction of that number (supplementary table S2). However, in both cases the stabilizing factor seems to be the electrostatic energies contributed by strong salt bridges and some ion-dipole interactions (Fig. 3d). Close inspection reveals that in our analyzed structures, there are a few similar salt-bridges formed. e.g. the *β*-catenin/*α*-catenin complex has an Arg90-Asp142 and an Arg93-Asp142 interaction at the interface (supplementary table S1), whereas *β*-catenin/IQGAP1 complex has a number of such salt-bridges, namely Arg90-Asp70, Arg93-Asp70, Arg124-Glu45, Arg124-Glu46 and HIS118-Glu46 (supplementary table S2). In addition, two types of *π* interactions (cation-*π* and *π*-H bond) seem to have reasonable contribution for these interfaces, as indicated by Fig. 3c and d. Therefore, the analyzed *β*-catenin/IQGAP1 interface appears to feature more favorable calculated pairwise electrostatic (ion-dipole and salt-bridge) interactions (Fig. 3d), compared to the analyzed *β*-catenin/*α*-catenin interface. Broadly, these provide a structural basis that can be used to understand the competitive binding of IQGAP1 and *α*-catenin with *β*-catenin (Kuroda et al. 1998; Fukata et al. 1999).

### 3.2. Effect of α-catenin Dissociation on IQGAP1 Docking

Our 100 ns MD simulation shows a very stable RMSD behavior of the *β*-catenin/*α*-catenin fragment complex with values between 0.40-0.42 nm and stable interfacial interactions. Reassured of the thermal stability of this complex, we proceed for its disruption by SMD. We simulate the force-dependent dissociation of the *α*-catenin fragment from a fixed *β*-catenin fragment using SMD at two pull velocities. Subsequently, we docked IQGAP1 to the *β*-catenin fragment of 15 chosen conformers from the SMD trajectories. The average docking scores for all the conformers, calculated from the scores observed across four replicates, are shown for these conformers in Fig. 4a. Here, scores are depicted for the partially blind (no active residue guidance) and the restrained protocols (active residue-guided). Scores from the completely blind protocol are presented in the supplementary Fig. S1. The interfacial area shared by *α*-catenin and *β*-catenin goes down for the conformers following a nearly sigmoid-like behavior (Fig. 4b) as the dissociation of *α*-catenin takes place. Three different conformers from a representative SMD trajectory are depicted in Fig. 4c-e sampled at different time points: initial (conformer 10 at 23 ps), intermediate (conformer 250 at 575 ps), and dissociated (conformer 500 at 1.15 ns), respectively. The best scoring IQGAP1 docked poses on these conformers from the residue-guided protocol are shown in Fig. 4f-h.

**Figure 4.:**
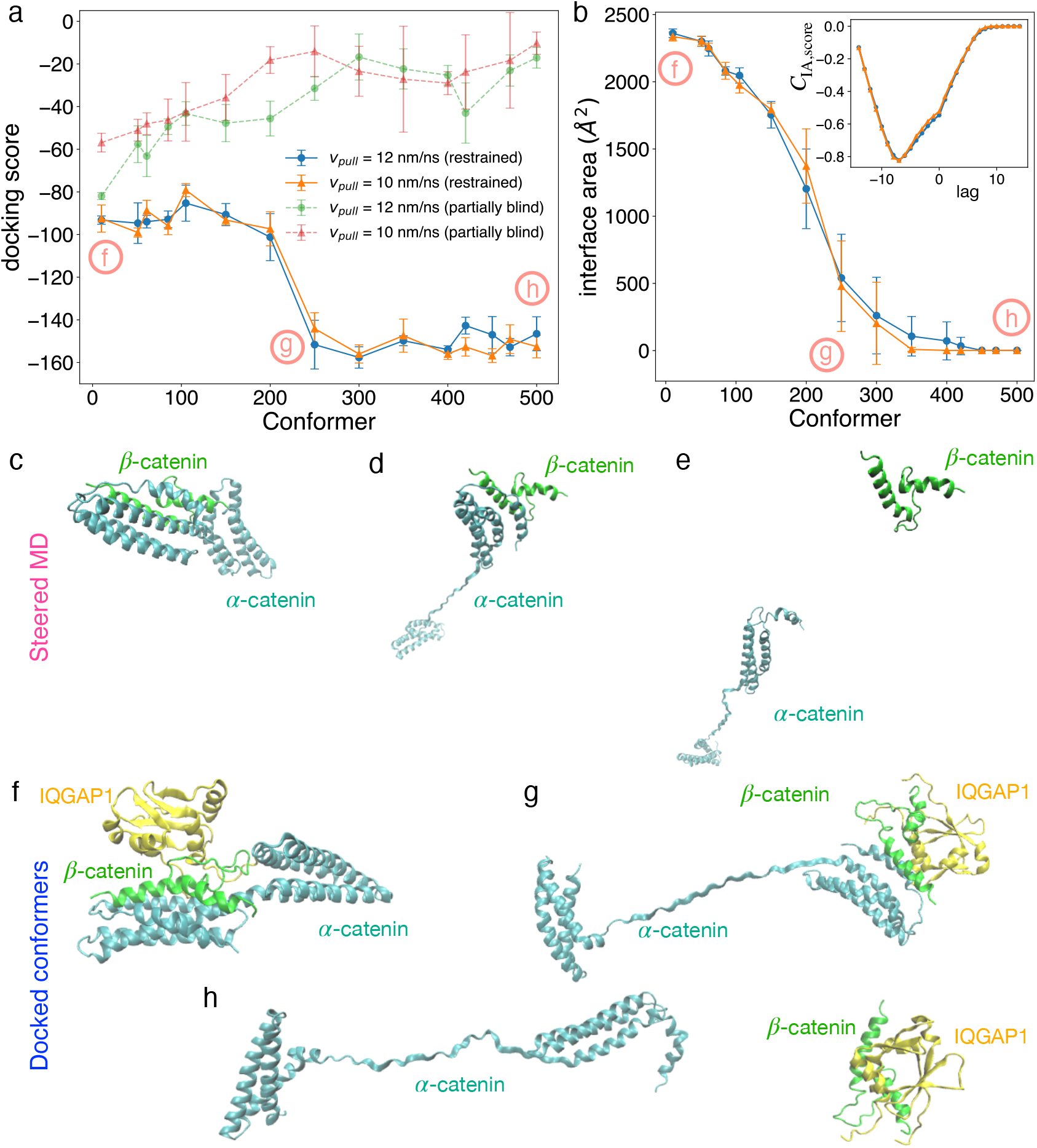
Results from SMD and step-wise docking protocol. (a) Docking scores for 15 conformers chosen from SMD trajectories generated for the *β*-catenin/*α*-catenin complex at two different pulling velocities. Data from two types of multi-step docking protocols are shown: partially blind, where docking of IQGAP1 is allowed anywhere on *β*-catenin, and restrained, where docking of IQGAP1 is done with specific active residues on *β*-catenin provided as AIRs. (b) Interfacial area of the *β*-catenin/*α*-catenin complex are shown for the 15 conformers from SMD trajectories. Here also data from two pulling velocities are shown. The interfacial areas are computed using PyMOL. Inset, results of cross-correlation analysis for the interface areas vs the docking scores. *C*_IA,score_ represents the normalized cross-correlation function, as a function of lag, measured in terms of conformer index (0 to 14) with interface areas taken as reference. All errorbars in panels a and b present standard errors of mean. (c) Three representative conformers from SMD simulations showing stepwise unbinding of *α*-catenin at different time points, like: (c)conformer 10 (23 ps), (d) conformer 250 (575 ps), and (e) conformer 500 (1150 ps). Lowest energy IQGAP1-docking poses for the (f) conformer 10, (g) conformer 250, and (h) conformer 500 are also shown. Here ribbon representations are shown in c-h where *β*-catenin is in green, *α*-catenin is in cyan and IQGAP1 is in yellow.

First we discuss the trends observed in the residue-guided case. Here, the docking scores demonstrate a clear trend that the docking of IQGAP1 becomes more favorable as *α*-catenin slowly detaches (Fig. 4a, b). Interestingly, we observe three clear stages. At early stages (conformer 10-200), the docking scores are insensitive to the state of detachment and extent to which this interface is disturbed. This is corroborated by the trends of docking scores and *β*-catenin/*α*-catenin interfacial area presented in Fig. 4a and b, respectively. The sharp increase in the magnitude of the negative docking score from conformer 200 to conformer 250 happens under conditions when the shared interfacial area between *α*-catenin and *β*-catenin is still significant, nearly 20-25% of that in conformer 10. This may be capturing the signature of initiation of some key stabilizing interactions offered by IQGAP1 to *β*-catenin. So, conformer 200 to 250 could represent a key stage in our multi-step docking. At late stages, conformer 250-500, the *β*-catenin/*α*-catenin interfacial area declines slowly (Fig. 4b), which most likely allows the IQGAP1 fragment to find the right binding pose. This is supported by the saturation of docking scores observed for conformer 250-500 where it fluctuates around an average value (Fig. 4a). All the above trends of docking scores remain consistent across all the replicates at both the pulling velocities (supplementary Fig. S2a).

In contrast, the above trends of docking scores divided into stages are totally absent in the scores obtained from the partially blind docking protocol, also shown in Fig. 4a. These average scores across the conformers exhibit little sensitivity to the continuous deformation of the *β*-catenin/*α*-catenin complex, with an overall weak, monotonically increasing trend. Such low sensitivity is expected, since there is no active residue guidance in this protocol, but binding anywhere on the surface of the *β*-catenin fragment is allowed. Interestingly, the saturated scores are slightly more favorable for the early stage conformers (*<* 250) compared to the latter ones (*≥* 250). We further assess these observations using interfacial analyses on the IQGAP1-docked poses of the conformers, which is described in the next subsection.

Next we check if above trends remain consistent in a completely blind docking protocol. Since this protocol allows IQGAP1 to explore the entire surface of the *β*-catenin/*α*-catenin complex for binding, we sometimes observe no interaction of the IQGAP1 fragment with *β*-catenin. Observed docking scores in this protocol, summarized in supplementary Fig. S1, shows insensitivity to the conformers.

Finally, a cross-correlation analysis is carried out to check if there is any delay between the progressive increase in favorable docking scores observed in the restrained protocol and continuous drop in interface areas. This allows us to assess whether the rising trend in the IQGAP1 docking score depends solely on the accessibility the *β*-catenin binding site. We consider the time-series of interface area as the reference, and evaluate its cross-correlation with the time-series of docking score, as a function of lag in terms of conformer index. These results for each of the four replicates are shown in Fig. S3. A negative peak value of the cross-correlation function (*C*_IA,score_), observed in case of each replicate, captures the opposing trends in the two quantities. Interestingly, this negative peak appears at a negative lag between the two time-profiles, which remains consistent across the replicates (Fig. S3a and b). We check the significance of this result by generating 100 random samples shuffling the docking scores from the 15 conformers. Then the mean lag observed from these 100 samples is compared with the originally observed negative lag using a Student’s t-test. The p-value observed for this test is *<* 10*^−^*^20^, which indicates that there is significant chance that the docking scores may be increasing at a rate little faster than the rate of decrease in the interface areas.

### 3.3. Role of electrostatic interactions

Our interfacial analysis reveals that both the protein-protein interfaces under consideration, namely *β*-catenin/*α*-catenin and IQGAP1/*β*-catenin, exhibit a dominant contribution from electrostatic interactions (Fig. 3c-d, and supplementary tables S1 and S2). Therefore, we focus on assessing the role of such interactions in a potential displacement of *α*-catenin by IQGAP1. First, we analyze all the salt-bridge interactions formed at IQGAP1/*β*-catenin interface in the docked conformers. Here we discuss the analysis done for SMD conformers chosen at pulling velocity 12 nm *ns^−^*^1^. We monitor the relative frequencies of occurrence of any *β*-catenin or IQGAP1 residue in a given docking model, given they are involved in a salt-bridge interaction at the docked interface. These relative frequencies, in case of each conformer, are calculated by pooling the interfacial data from the top ten docking models observed for each of the four replicates, *i.e.* a total of 40 models. These results are summarized for six conformers, in Fig. 5, for the partially blind docking protocol (Fig. 5a,b), the restrained protocol (Fig. 5c,d), and in supplementary Fig. S4 for the fully blind protocol.

**Figure 5.:**
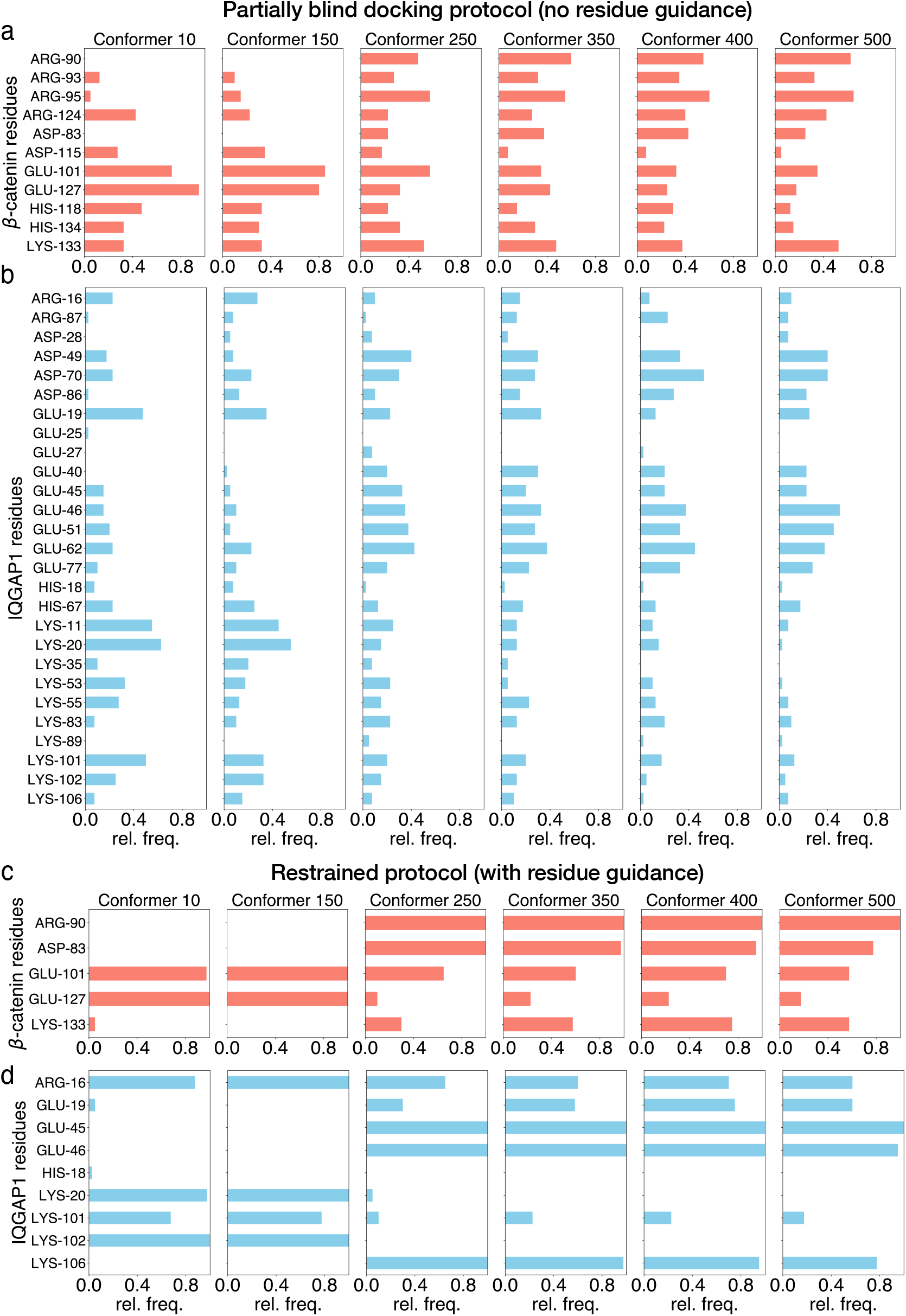
Results of interfacial analysis on IQGAP1/*β*-catenin interface in six different conformers. Data are shown from two different docking protocols: (a-b) Partially blind, with no residue guidance and (c-d) restrained, with active residue guidance. Here, the relative frequencies of occurrence of all *β*-catenin residues (panels a and c) and IQGAP1 residues (panels b and d), involved in any salt-bridge interactions estimated with a 5 Å distance cutoff, are shown in form of bar graphs. Here we have used a larger distance cutoff for better statistics. The relative frequencies are computed from the top ten docking models for each conformer across four independent replicates. When residues are recovered in multiple salt-bridges in a given model, they are considered only once. So, a relative frequency of 0.5 indicates that the corresponding residue is recovered in at least one salt-bridge in 50% of 10 *×* 4 structures analyzed.

The trends from Fig. 5 indicate that the partially blind protocol involves many more residues forming salt-bridges between the two fragments compared to the restrained protocol. This is expected from the completely relaxed AIRs implemented in the former protocol. However, some of the residues involved in the later stage conformers (*≥* 250) are also recovered in both the protocols. E.g. Arg90, Asp83, Glu101, Glu127 and Lys133 from the *β*-catenin fragment and Arg16, Glu19, Glu45, Glu46, Lys101 and Lys106 from the IQGAP1 fragment. On the other hand, some of the residues with high occurrence frequency during the partially blind protocol are not recovered in the restrained protocol, like Arg95, Arg124 (*β*-catenin fragment) and Asp49, Asp70, Glu51, Glu62 and Glu77 (IQGAP1 fragment). Similarly, some of the IQGAP1 residues, like Arg16, and Lys106 show high occurrence frequency in the restrained docking models only, while the presence of others like Glu19, Glu45 and Glu46 remain reasonable to high across the protocols. In comparison, these overall occurrence frequencies in the fully blind protocol become at least ten times lower than those observed in the other two. This reflects the completely blind nature of the protocol. Close inspection shows that the IQGAP1 fragment sometimes binds even to the *α*-catenin fragment, missing the *β*-catenin fragment completely as indicated by the docked structures shown in Fig. S1. However, even with these relaxations, most of the *β*-catenin as well as several of the IQGAP1 residues recovered in the restrained model are also observed in the fully blind model across the conformers. The overall findings and comparative behaviors of the three protocols remain consistent across the pulling velocities, so we concentrate only on a single velocity as indicated above.

Next, we analyze the specific residue-level interactions for docked conformers with favorable docking scores. For this we centre our discussions on the restrained protocol as this is the only protocol among the 3 that shows favorable docking scores as *α*-catenin detaches from *β*-catenin. Since all replicates in this protocol show qualitatively similar variation of docking scores at the two pulling velocities (Fig. S2a and b), going forward we only focus on the conformers chosen from the SMD trajectories at pulling velocity 12 nm *ns^−^*^1^. Here, we only show data from replicate 1.

Table 1 enumerates the interfacial residues and electrostatic contacts observed at the IQGAP1/*β*-catenin interface in all the three conformers shown in Fig.4f-h. Conformer 500, representing the fully dissociated state of *α*-catenin from *β*-catenin, is among the best scoring docked conformers (Fig. 4a). Here, we observe an significant number of assorted electrostatic interactions (Fig. 6a-c), including 4 different salt bridges, 7 ion-dipole, 6 H-bonds, and 9 different interactions involving aromatic side-chains with ionic (cation-*π*) or dipolar side-chains (*π*-H bonds). A few of these specific contacts do form in some of the conformers at the early stages (conformer 10-200), as shown by the interfacial residues shown in Table 1, and supplementary Table S4 (conformer 150), S5 (conformer 200) and S7 (other replicates). Several of these specific interactions could be observed in conformer 250 and following late-stage conformers (Fig. 5c-d, supplementary Table S6 and S7).

**Table 1.:**
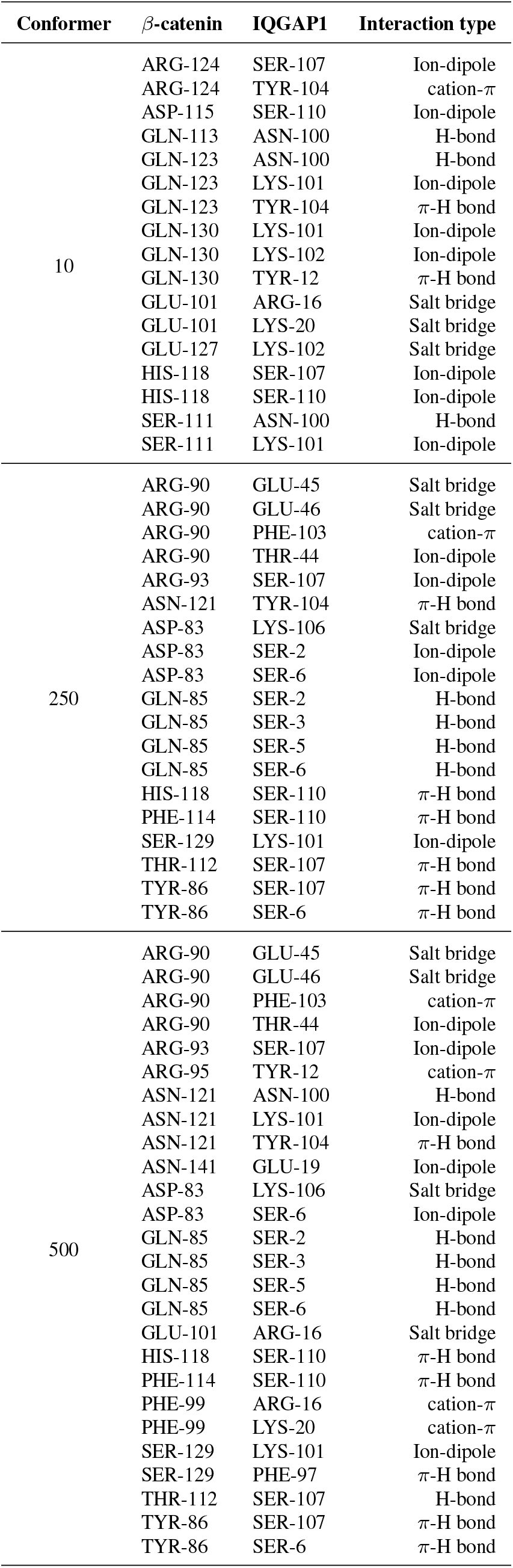
Interfacial electrostatic contacts observed for the docked IQGAP1 and the conformers chosen from SMD.

**Figure 6.:**
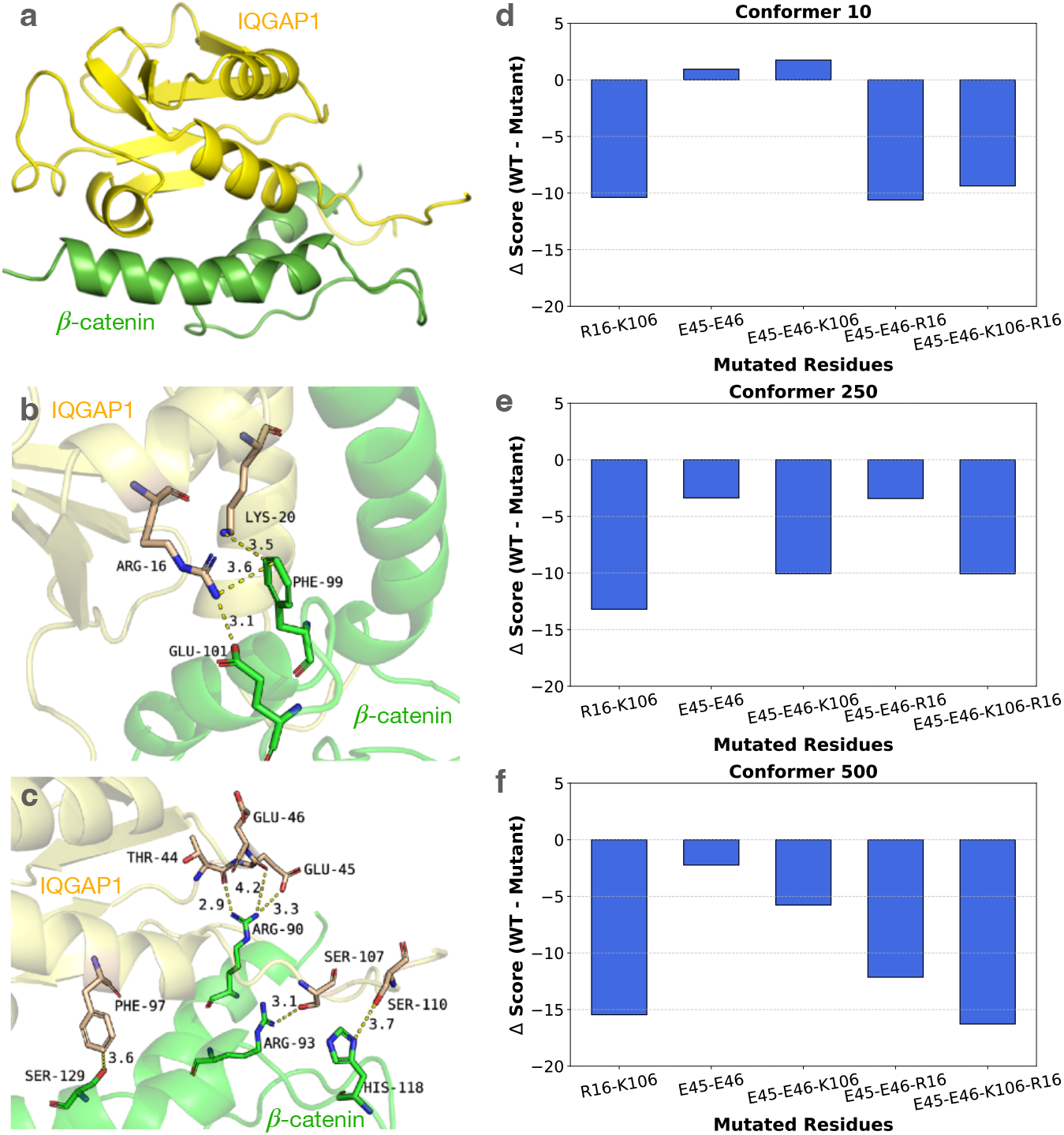
Importance of ionic and polar interfacial interactions in stabilizing IQGAP1-*β*-catenin complex. (a) IQGAP1 docked with conformer 500. (b-c) Zoomed view of the interface of IQGAP1-conformer 500 complex with key salt-bridges, ion-dipole contacts and H-bonds. (d-f) Comparison of Δ-scores (WT score *−* mutant score) for several IQGAP1 mutants. Here we mutated several key interfacial residues of IQGAP1 to prepare double, triple and quadruple mutants. Docking scores are shown for IQGAP1 mutants in 3 conformers: conformer 10 (d), conformer 250 (e) and conformer 500 (f).

Subsequently, we use conformer 500 to test the hypothesis that specific electrostatic interactions could be key to IQGAP1 binding to *β*-catenin. For this we target the four salt bridges: Arg90-Glu45, Arg90-Glu46, Asp83-Lys106 and Glu101-Arg16. All these contacts are recovered consistently in the restrained protocol across replicates for the late-stage conformers (*≥* 250). The three *β*-catenin residues involved are consistently recovered with moderate to high occurrence frequencies across protocols (Fig. 5a and c). Among the IQGAP1 residues involved, Glu45 and Glu46 are consistently recovered across protocols with moderate to high occurrence frequencies, while Arg16 and Lys106 are also recovered across protocols but with low occurrence frequencies (Fig. 5b and d).

We decide to disrupt these interactions by mutating all four of the involved residues on IQGAP1 to alanine directly in the binding interface of IQGAP1 present in the docked conformers. In addition to this quadruple mutant, we also check the scores of the double mutants (Glu45-Glu46 both mutated to Ala and Arg16-Lys106 both changed to Ala) and triple mutants (Glu45-Glu46-Arg16 mutated to Ala and Glu45-Glu46-Lys106 mutated to Ala). Our recalculated docking scores for all the mutant IQGAP1 on the three different SMD conformers are summarized in Fig.6d-f in form of a Δ-Score (Wild type score - mutant score). A negative Δ-Score is indicative of a poorer docking at the mutant IQGAP1/*β*-catenin interface compared to the corresponding wild-type case. The quadruple mutant results in the biggest drop in docking score, with the Δ-Score ranging in *−*10 (conformer 10) to *−*17 (conformer 500) (Figure 6d-f). This drop is also consistent with an enhanced RMSD and reduced interfacial area in the quadruple mutant as revealed by our subsequent MD simulations (supplementary Fig. S5 and S6). Above observations indicate that these residues could be good candidates for setting up favorable salt-bridges at the *β*-catenin/IQGAP1 interface. Interestingly, 3 out of 4 of these salt bridges move into interacting position quite early in our serial docking simulations, among other salt-bridge pairings (see Table 1 and Fig. 5d). This evidence highlights the importance of favorable salt-bridge interactions in the proposed mechanism of IQGAP1-driven disruption of the *β*-catenin/*α*-catenin complex.

### 3.4. RMSD and interface area comparison between wild-type and mutated IQGAP1 fragments bound to β-catenin

Next, we compare the RMSD as well as interface areas of the wild-type mutant IQGAP1/*β*-catenin fragment complexes estimated from the MD trajectories. We consider data from four independent replicates for each case. This allows us to assess the structural stability and the extents of interfacial overlap of these complexes along the MD trajectories. Two types of RMSD are considered: the RMSD of the protein backbone (Fig. S5a), which primarily captures the stability of the protein secondary and tertiary structures; and the RMSD of the whole protein (Fig. S5b), which captures the fluctuation of the overall protein structure including those of the side-chains. Both these types of RMSD are close in magnitude and temporal fluctuation for a given complex and capture same trends. RMSD for the wild-type complex and the two double mutants remain between 0.3-0.7 nm throughout the simulations (Fig. S5a-b, i-iii), indicating a stable structure and moderate conformational fluctuation for these 3 protein complexes. In contrast, the RMSD of *β*-catenin docked with the quadruply mutated IQGAP1 fragment (Arg16-Glu45-Glu46-Lys106 all mutated to ALA) increases to approximately 0.8 nm with significantly enhanced variations over the trajectories (Fig. S5a-iv and b-iv), suggesting much higher structural fluctuations and possibly reduced complex stability. The temporal profiles of interface area capture similar behavior as the RMSD (Fig. S6a).

In this case, we also include the results of unpaired significance tests involving every pair of timeseries shown in Fig. S6b. These results suggest that the interfacial area in the Arg16Ala-Lys106Ala complex maybe significantly lowered because of the mutations, while the Glu45Ala-Glu46Ala-mutations seem to have no significant effect. The quadruple-mutant clearly shows significantly lowered interfacial area among all 4. Visual inspection of the simulation snapshots indicates reasonable deformation as the IQGAP1 fragment partially detaches from *β*-catenin. This explains the observed reduction in interface area and large variations across the replicates. These trends with quadruple mutant IQGAP1, along with large variations, is consistent with the disruption of favorable electrostatic interactions.

## 4. Discussion and Conclusion

One of the main ingredient of cell–cell junctions are the adhesion complexes, which are subjected to dynamic exchange with their cytoplasmic populations (Founounou et al. 2021; Iyer et al. 2019) or remodeling under mechanical forces (Curran et al. 2017). Often the adhesions can be disrupted by various mechanical factors, like, abnormal build up of myosin-driven contractile forces (Li et al. 2021; Duque et al. 2024), external shear/injury (Colpitts and Grau 2026) or cell-matrix interactions via focal adhesions (Wu et al. 2026). Apart from mechanical means, biochemical routes for the disruption has also been observed before in form of IQGAP1-driven dissociation of *α*-catenin from the essential adhesion complex of E-cadherin-based cell-cell junctions (Fukata et al. 1999; Kuroda et al. 1998). When IQGAP1 is over-expressed, as is the case in various disease conditions, it is known to associate with *β*-catenin (Jin et al. 2015; Hu et al. 2019), a key part of the adhesion complex with *α*-catenin. According to previous reports (Fukata et al. 1999; Kuroda et al. 1998), a competitive binding of IQGAP1 and *α*-catenin on an overlapping binding site on *β*-catenin has been implicated in this process. However, any relevant molecular details regarding a key binding interface of IQGAP1 has been missing. Here, we employ a multi-step docking of IQGAP1 on a whole spectrum of fully bound, partially bound and completely unbound *β*-catenin/*α*-catenin complexes to predict a molecular basis for this disruption and a set of candidate residues of IQGAP1, which may assist the process.

Our regular equilibrium MD simulations show that the *β*-catenin/*α*-catenin complex remains stable under unperturbed conditions, with favorable electrostatic interactions and hydrophobic contacts. When we induce a detachment of *α*-catenin by mechanical pulling, it becomes extended in distinct steps with unfolding of specific helical domains. Such open conformation of *α*-catenin has been alluded to be a key intermediate step of cytoskeleton-driven regulation of the junctional strength (Buckley et al. 2014; Adhikari et al. 2018; Li et al. 2015; Barrick et al. 2018). This forced detachment of *α*-catenin allows us to tune the accessibility of the *β*-catenin interface for binding of IQGAP1. Stepwise docking shows that the docking of IQGAP1 to *β*-catenin in the deformed complex becomes favorable in a progressive manner (Fig. 4a). This could be attributed to the increasing exposure of the residues in the overlapping binding site of *β*-catenin that have been interacting with *α*-catenin residues. A continuous drop in interfacial areas across the progressively deforming conformers suggests so. But there could be more than just the factor of increasing exposure. Our analysis shows a negative cross-correlation between the drop in interfacial area and the docking scores (Fig. 4b, inset), which indicates a faster increase in favorable docking scores than that expected from the increased exposure of *β*-catenin binding site due to the detachment of *α*-catenin. Subsequently, our detailed comparison of the two binding interfaces in *β*-catenin/*α*-catenin complex and *β*-catenin/IQGAP1 complex (wild type as well as exposed *β*-catenin, like conformer 250-500) reveals that the latter is able to support favorable salt-bridges and other types of electrostatic interactions. These different pieces of evidence taken together support a probable molecular mechanism for IQGAP1 binding to a mechanically weakened *β*-catenin/*α*-catenin complex.

Our stepwise docking workflow also supports a dynamic picture of mechanical modulation of a protein-protein complex and provides a stepwise illustration of how mechanochemical coupling can occur at the molecular scale. Inspection of the interfacial residues in the SMD conformers reveals an early engagement of some of the key interacting IQGAP1/*β*-catenin residue-pairs (Table 1, Table S4, S5 and S7), while *α*-catenin is still partially bound to *β*-catenin. These engagements are seen as early as conformer 200 to 250, the exact middle point of unbinding. Therefore, less than 50% detachment of *α*-catenin from the adhesion module could be enough for IQGAP1 to form its important interactions as predicted in this work. The fact that the docking score profile appears to ‘lead’ the interfacial area drop supports this suggestion. Such partial detachment situations could be physiologically relevant because of recent report (Windgasse and Grashoff 2025) that *α*-catenin, at mature cell-cell junctions, undergo dynamic conformation switching and enhanced turnover. Future single-molecule experimental studies using optical tweezers and super-resolution imaging could test the proposed molecular mechanism for IQGAP1 binding to the partially-deformed states of the adhesion complex and possibly validate some of our predicted molecular details.

Interfacial analyses further highlight Glu45, Glu46, Arg16 and Lys 106 on IQGAP1 as important candidate residues emerging through comparative docking protocols, which could form stabilizing salt bridges with *β*-catenin. Mutating all four of these residues to alanine significantly reduces the docking score for IQGAP1. Subsequent MD simulations of the resultant complex show reduction in interfacial area between mutated IQGAP1 and *β*-catenin (Fig. S6), and also substantially enhanced RMSD values (Fig. S5). These pieces of evidence give us some clue for screening a few small molecules that can target this binding interface of IQGAP1 and inhibit its disruption activity. Our investigation of existing literature shows two previously suggested inhibitors of IQGAP1 (NCBI 2026b,a). Our preliminary docking shows at least one of them is able to interact with three of the above-mentioned charged residues on our predicted binding interface of IQGAP1. These docking results are summarized in supplementary Fig. S7. This evidence may be considered for generating hypothesis for a potential design of an IQGAP1-inhibitor preventing its competitive binding to *β*-catenin.

Although our framework predicts a molecular mechanism for how IQGAP1 could bind to partially-deformed to fully-deformed adhesion complex, along with a number of candidate residues for favorable binding, our approach remains limited by the absence of full-length versions of the binding proteins involved. So, we may be missing a large amount of information related to residues involved in binding and potential hotspots for targeting. Furthermore, we could not capture the cytoplasmic crowding effect on the proteins, which is expected to significantly influence the quantitative details of the proposed mechanism. All the three proteins involved have other binding partners known to exist near the junctional area, *e.g.* E-cadherin cytoplasmic domain for *β*-catenin, and vinculin, afadin for *α*-catenin (Terekhova et al. 2019). *α*-catenin also forms dimers and oligomers, which are essential for maturation of adherens junctions (Troyanovsky, Indra, and Troyanovsky 2025). Active Rac1 and Cdc42 are known to be regulators of IQGAP1 activity (Owen et al. 2008; White, Erdemir, and Sacks 2012; Nouri, Timson, and Ahmadian 2020). Finally, IQGAP1 undergoes a series of post-translational modifications, namely phosphorylation of several serines and tyrosines (Table 1) (Hedman et al. 2020) and ubiquitination (Gorisse et al. 2020) that also regulate its activity. So, a comprehensive study of this problem would require a consideration of these various activation and inactivation scenarios of the central proteins. Another potential limitation of our study is that the ionic strength used is on the lower end of the physiological range of 0.1-0.3 M (Dzubiella 2008). Higher values of ionic strength at around pH=7-8, compared to which which we have used, has been shown to reduce the overall stability of protein-protein interactions to some extent (Roberts et al. 2015). Sensitivity of biomolecular interactions to specific cationic species is also another important factor (Okur et al. 2017). We use Na^+^ ions, while the dominant cationic species inside the cell is K^+^. This can have ion-specific influence on the stability of the important salt-bridge interactions as Na^+^ has been implicated (Vrbka et al. 2006) to have higher binding affinity to protein surfaces compared to K^+^ and hence may destabilize electrostatic interactions among protein side-chains. These disruptions could be potentially more effective for solvent-exposed salt-bridges compared to buried ones as suggested earlier (Luo et al. 1999). However, nearly all recent work (Okur et al. 2017; Taghavi Shahraki and Khajeh-pour 2025) have suggested that predominantly at much higher salt concentrations (Dzubiella 2008) (*>* 1 M or so) any destabilizing effect of the alkali monovalent cations are realized. More importantly, at the salt concentrations we have used, ion-specific differences between Na^+^ and K^+^ in context to stability of biomolecular interactions remain negligibly small (Okur et al. 2017).

Finally, an ideal protocol for probing the competitive binding would be needed to simulate an in situ docking of IQGAP1 during the simulation of forced detachment of *α*-catenin. That is technically very hard to achieve as the MD methods remain limited at simulating binding-unbinding events. Here we have tried an alternative – simulate the forced-detachment, sample conformers at various stages of detachment and perform a multi-step docking on the partially detached complex using these sampled conformers. Such a protocol is definitely limited by a natural obstacle as the binding site for IQGAP1 remains occupied by *α*-catenin. Interestingly, the restrained docking protocol shows that even before 50% of detachment of *α*-catenin (around conformer 200, as shown by interface data in Fig. 4b), IQGAP1 is able to engage some of its favorable electrostatic interactions with *β*-catenin (Table S5 and S7). Our result related to the negative cross-correlation between the drop in interfacial areas and increase in favorable docking scores appear to provide support for this favorable effect (Fig. 4b, inset), which could lead to future investigations.

In conclusion, our findings suggest that mechanically weakened adhesion complex at cadherin-catenin junctions may have higher chances of getting disrupted by IQGAP1. Our multi-step docking does not directly address force-dependent exchange. However, the results related to increase in favorable docking scores as detachment happens and early onset of consistent salt-bridges between IQGAP1 and *β*-catenin even when the detachment of *α*-catenin is largely incomplete, indirectly point to a probable molecular mechanism for this disruption process. In addition, the computational approach we adopt here, linking MD, SMD, stepwise docking, interface analysis and *in silico* mutagenesis, offers a strong and adaptable framework for exploring other mechanically regulated protein–protein interactions (Amieva et al. 2003; Romero-Fabela et al. 2025) that are crucial to cellular mechanobiology.

## 5. Data availability statement

The authors confirm that most of the data supporting the findings of this study are available within the article and its supplementary materials. Any additional data and the analysis and simulation codes are available from the corresponding author, Amit Das, upon reasonable request.

## 6. Declaration of generative AI and AI-assisted technologies in the manuscript preparation process

During the preparation of this work the author(s) used Gemini Pro and ChatGPT in order to rephrase some parts of the text and suggestions of complex literature on specific topics. After using this tool/service, the author(s) reviewed and edited the content as needed and take(s) full responsibility for the content of the published article.

## Credits

### 6.0.0.1. Author Contributions

NB: Investigation, Formal analysis, Writing – Original Draft and editing. AD: Conceptualization, Funding acquisition, Investigation, Formal analysis, Editing the Draft.

### 6.0.0.2. Funding

AD acknowledges funding support from the New Faculty SEED Grant from IIT Delhi, India and the Start-up Research Grant from ANRF (previously SERB-DST), Government of India (Grant No. SRG/2023/000099 and ANRF/ARG/2025/003382/LS). Authors acknowledge support from IIT Delhi High Performance Computing facilities.

