## Supplementary Material for "Understanding the role of IQ motif-containing GTPase-activating protein 1 as a potential disruptor of the essential adhesion module at cell-cell junctions"

### Supplementary Text

#### Estimation interfacial interaction energies using INTAA webserver

Interaction energies were obtained using the INTAA webserver (Vymětal et al. 2021) using the following technique: The structures (4ONS for $\beta$-catenin-$\alpha$-catenin complex, and docked $\beta$-catenin-IQGAP1 structure) were uploaded after removing the H-atoms. The webserver application was executed with the 'add hydrogens' mode. The OPLS-AA force-field was chosen for the energy calculations. The output from the webserver was processed with in-house python script. We filtered the output distances matrix for all side-chain-side-chain separations $\leq$ 4 Å. The residue-residue interaction energies for all detected residue-pairs satisfying this distance criteria were considered for reporting in the tables below.

### Supplementary Figures

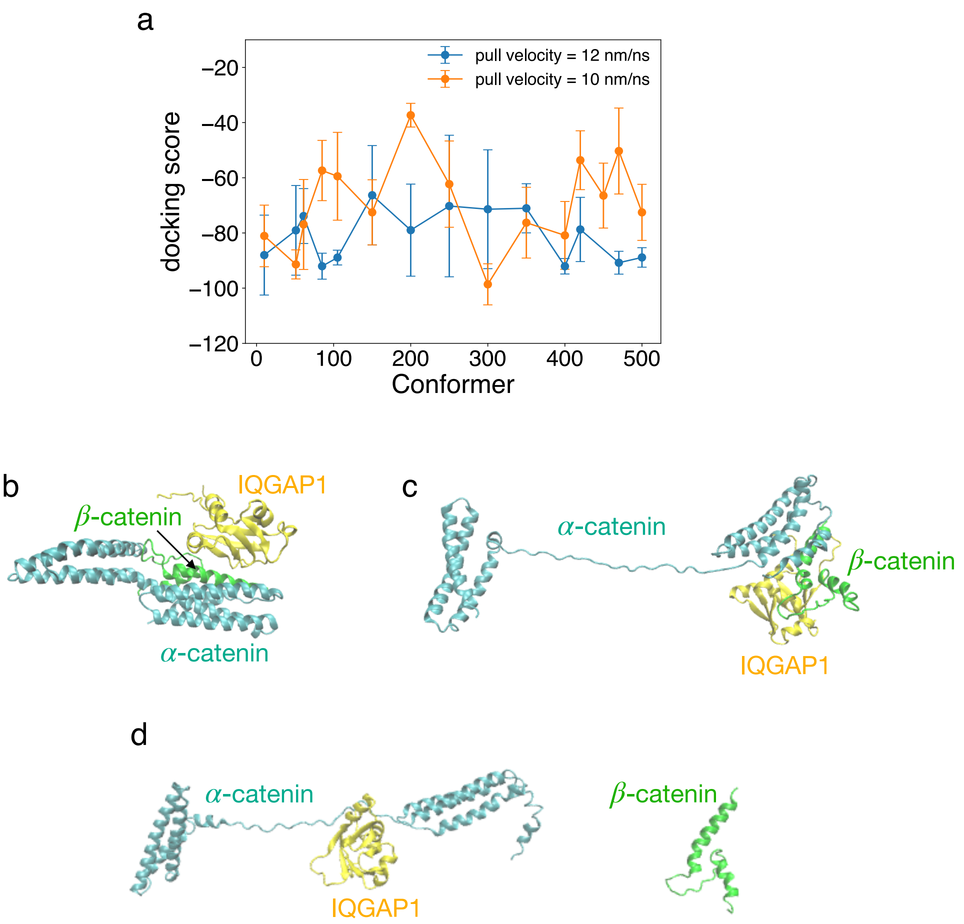


Figure S1: (a) HADDOCK scores for completely blind docking of IQGAP1 on 15 conformers chosen from SMD trajectories generated for the $\beta$-catenin/$\alpha$-catenin complex at two different pulling velocities. Here any type of residue guidance is removed and also spatial restraints are also removed. (b-d) Representative docked poses observed in this protocol. Here the conformers involved are shown in main figure 4c-e. Because of the fully-blind nature of this protocol, sometimes the IQGAP1 fragment docks completely on the $\alpha$-catenin fragment as indicated by panel d. Subsequently, these configurations are not included for additional interaction analyses.

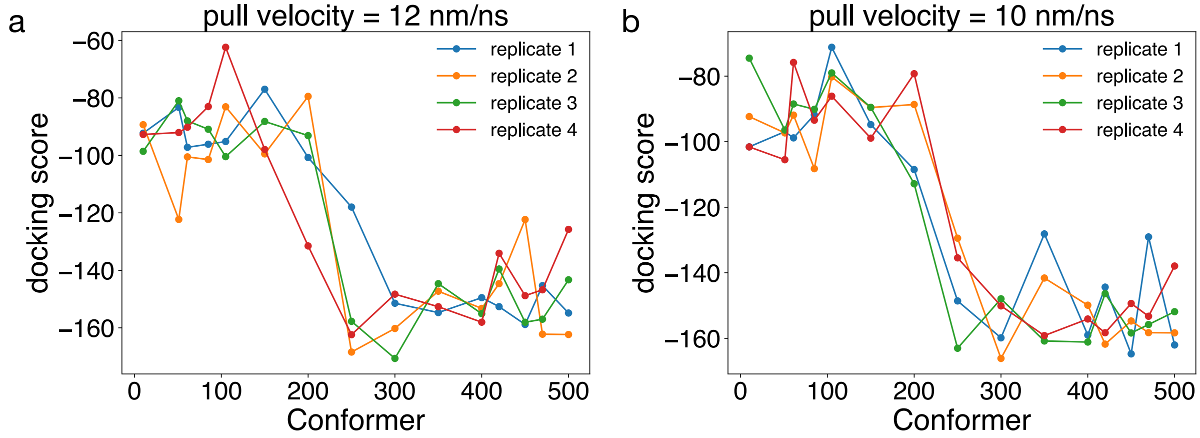


Figure S2: Raw docking scores observed for all the replicates 1-4 for pull velocities (a) 12 and (b) 10 nm/ns. These replicates are from the active residue guided restrained protocol.

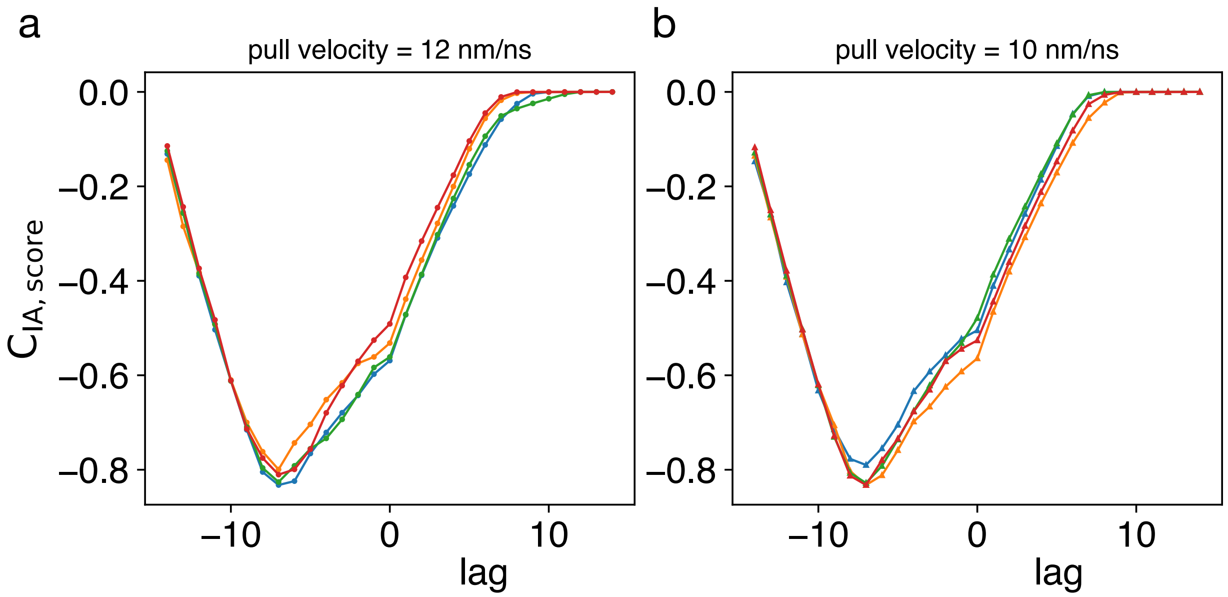


Figure S3: Replicate-wise results of cross-correlation analysis for the interface area profile vs the docking score profile, observed in the restrained protocol, at pull velocities (a) 12 and (b) 10 nm/ns. Each line shown represents the normalized cross-correlation function, $C_{\mathrm{IA,score}}$ as a function of lag, measured in terms of conformer index (0 to 14). Here, the cross-correlations are calculated with interface areas taken as the reference signal.

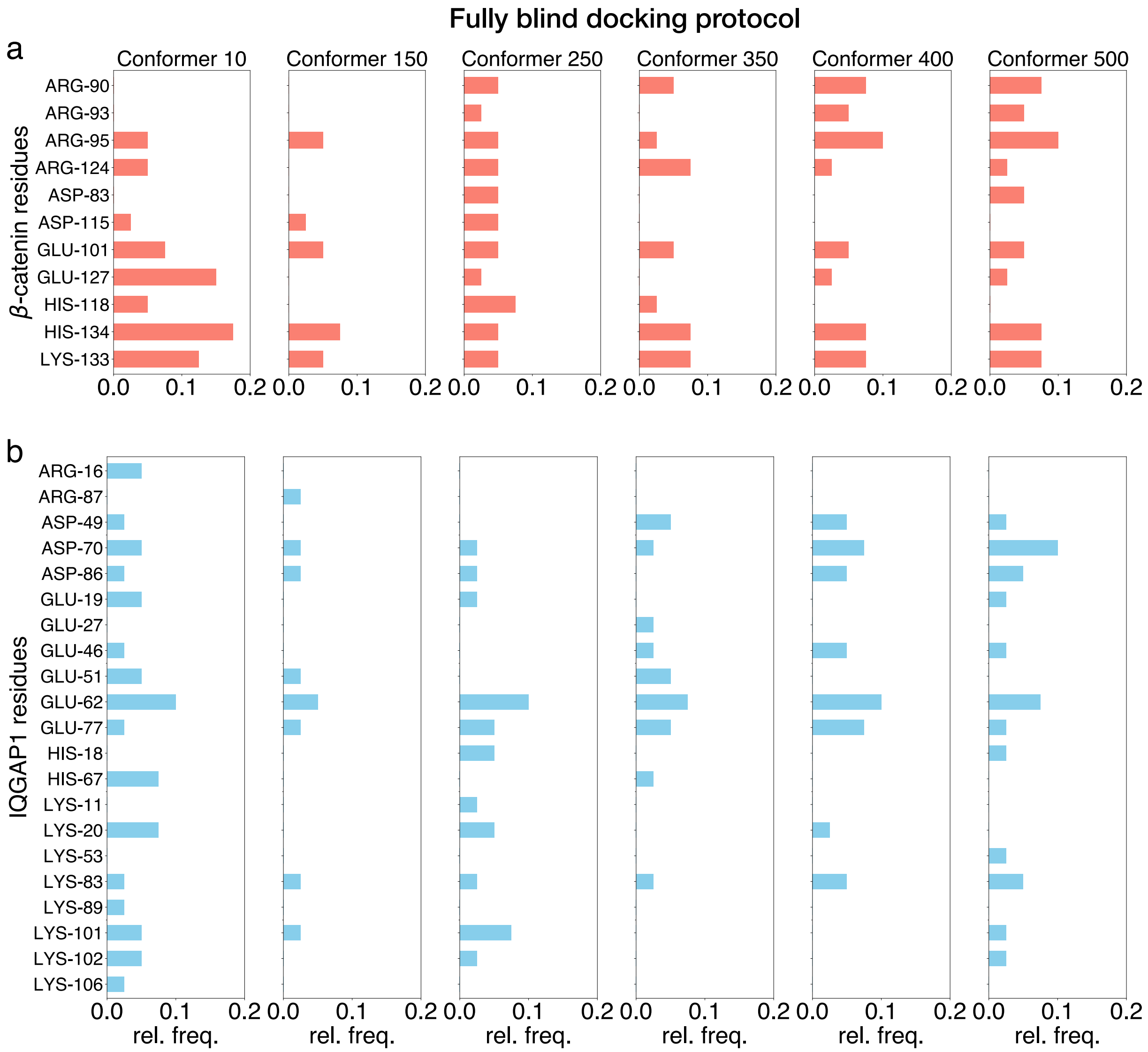


Figure S4: Results of interfacial analysis on IQGAP1/$\beta$-catenin interface in six different conformers from the completely blind docking protocol: Here, the relative frequencies of occurrence of all $\beta$-catenin residues (panels a) and IQGAP1 residues (panels b), involved in any salt-bridge interactions, detected with a 5 Å distance cutoff, are shown in form of bar graphs. The relative frequencies are computed from the top ten docking models for each conformer across four independent replicates. So, relative frequency of 0.5 indicates that the corresponding residue is recovered in 50% of $10 \times 4$ structures analyzed.

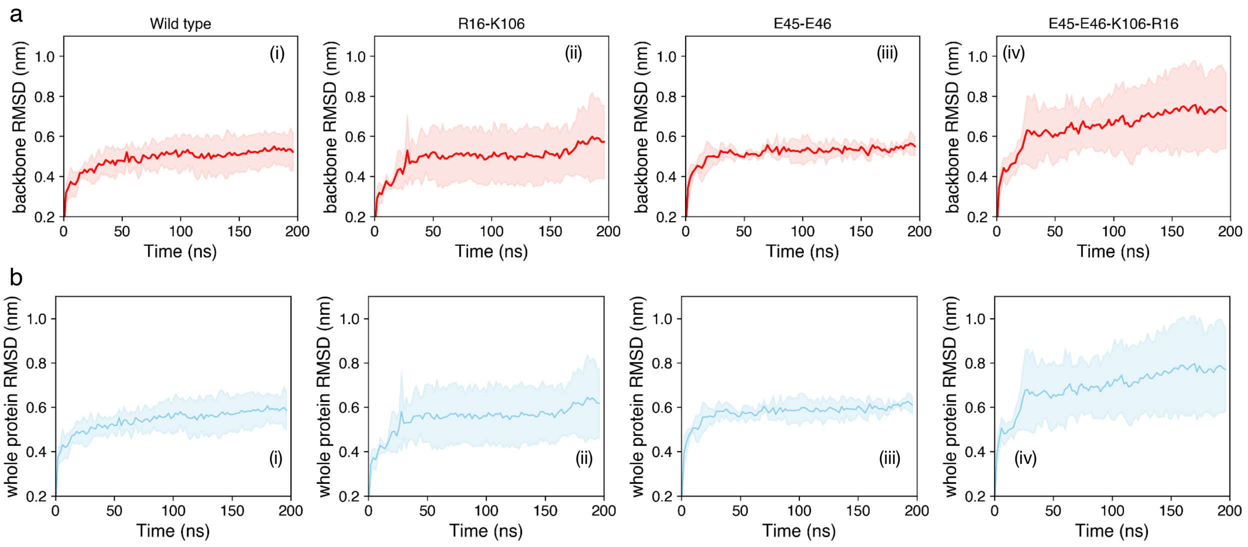


Figure S5: Overall structural fluctuations of IQGAP1-$\beta$-catenin complexes during MD runs in form of Root mean square deviations (RMSD) of (a) the protein backbone and (b) the whole protein, for four different IQGAP1 fragment complexes with $\beta$-catenin: (i) wild type IQGAP1 fragment, (ii) IQGAP1 double mutant Arg16Ala-Lys106Ala, (iii) IQGAP1 double mutant Glu45Ala-Glu46Ala, and (iv) IQGAP1 quadruple mutant Arg16Ala-Lys106Ala-Glu45Ala-Glu46Ala. In each case, the solid line represents the average from four replicates, while the shaded region represents the standard deviation observed at each time-point across the replicates.

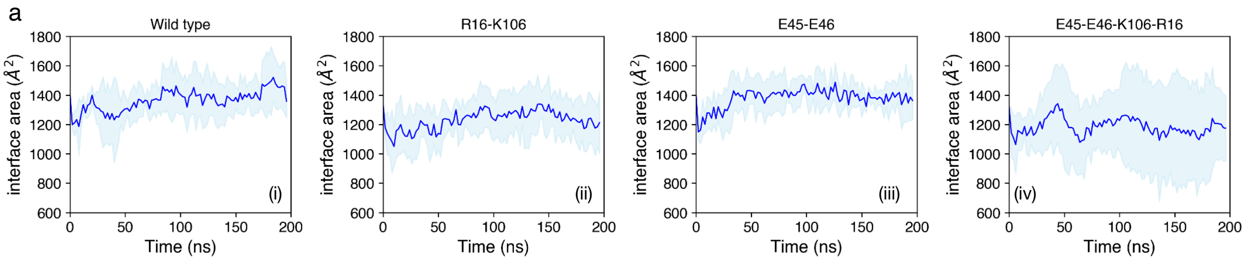


b

Significance test results of interface area

| sample1 | sample2 | p-value |
|---|---|---|
| Wild type | R16-K106 | $< 10^{-10}$ |
| R16-K106 | E45-E46 | $< 10^{-10}$ |
| Wild type | E45-E46 | $> 0.05$ |
| R16-K106 | E45-E46-K106-R16 | $< 10^{-7}$ |
| E45-E46 | E45-E46-K106-R16 | $< 10^{-10}$ |
| Wild type | E45-E46-K106-R16 | $< 10^{-10}$ |

Figure S6: (a) Interface area variation during MD runs of four different IQGAP1 fragment complexes with $\beta$-catenin: (i) wild type IQGAP1 fragment, (ii) IQGAP1 double mutant Arg16Ala-Lys106Ala, (iii) IQGAP1 double mutant Glu45Ala-Glu46Ala, and (iv) IQGAP1 quadruple mutant Arg16Ala-Lys106Ala-Glu45Ala-Glu46Ala. In each case, the solid line represents the average from four replicates, while the shaded region represents the standard deviation at each time-point across the replicates. (b) Results of significance tests for various pairs of samples are shown. For each pair mentioned as sample 1 and sample 2, an unpaired independent sample Student's t-test is used.

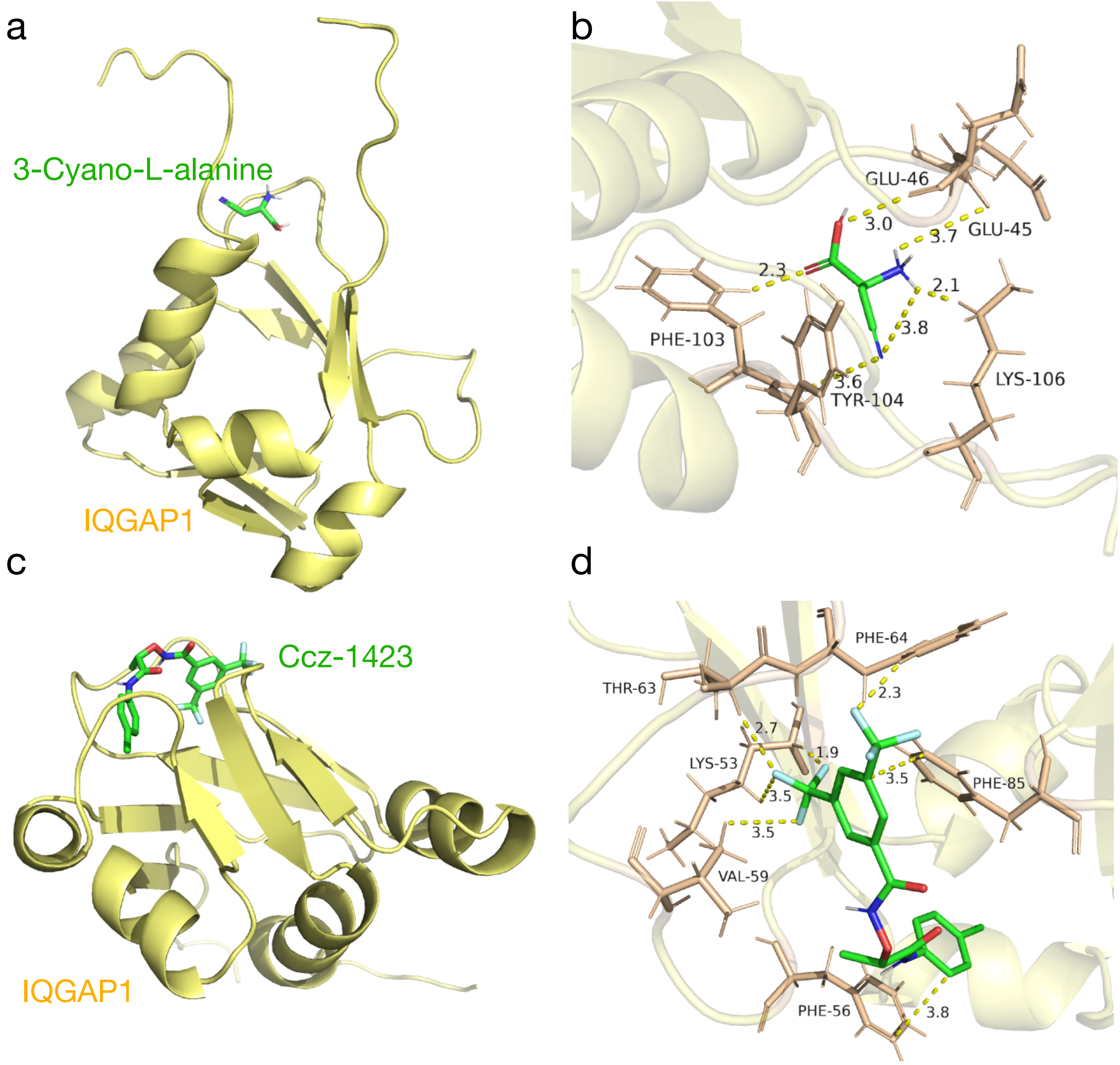


Figure S7: Two small molecules docked on IQGAP1 fragment: (a-b) 3-Cyano-L-alanine and (c-d) Ccz-1423 (IUPAC name: N-(1-((4-chlorophenyl)amino)-1-oxopropan-2-yl)oxy-3,5-bis(trifluoromethyl)benzamide). We used AutoDock Vina (Trott and Olson 2010; Eberhardt et al. 2021) through UCSF Chimera (Pettersen et al. 2004) to perform the docking. Panels a and c show a full view of the protein fragment with the given small molecule and panels c and d show a zoomed-in view of the various interactions of the given small molecule with different protein residues. Several close contacts $\leq 4.0$Å are shown as dashed lines with distances mentioned in Å. The two small molecules show different docking sites in the all the scoring models. 3-Cyano-L-alanine (best docking score $-4.4$) interestingly binds to a region where it makes various close contacts below 4.0 Å with Glu45, Glu46 and Lys106, which are proximal residues. On the other hand, Ccz-1423 shows a much higher docking score of $-8.4$ but docks in a different site where it makes contact with several phenylealanine residues (Phe-56, 64 and 85), Lys53 and Thr63. Given its significant aromatic content this is expected. All panels here have been prepared with PyMOL and distances are calculated using the ‘Measure Distances’ wizard in PyMOL.

#### Supplementary Tables

Table S1: **Residue-level interaction analysis of $\beta$-catenin-$\alpha$-catenin complex**

| $\beta$-catenin Res | $\alpha$-catenin Res | Distances (Å) | Energies (KJ/mol) | Interaction type |
|---|---|---|---|---|
| ARG-90 | ASP-142 | 2.257 | -80.218 | Salt bridge |
| ARG-93 | ASP-142 | 1.752 | -181.832 | Salt bridge |
| ARG-124 | GLU-81 | 2.082 | -74.338 | Salt bridge |
| ASP-144 | LYS-56 | 2.556 | -28.834 | Salt bridge |
| TYR-142 | PHE-110 | 2.151 | -7.046 | $\pi-\pi$ |
| GLN-113 | TYR-188 | 3.49 | -5.919 | $\pi$-H-bond |
| TYR-142 | SER-54 | 1.481 | 57.428 | $\pi$-H-bond |
| GLN-85 | ARG-193 | 3.547 | 0.484 | ion-dipole |
| SER-111 | ARG-192 | 2.846 | 4.684 | ion-dipole |
| THR-112 | ASP-145 | 2.261 | 3.148 | ion-dipole |
| GLN-113 | ARG-192 | 3.878 | -1.507 | ion-dipole |
| THR-120 | ASP-145 | 2.316 | -47.26 | ion-dipole |
| ASN-121 | ASP-145 | 2.55 | -80.322 | ion-dipole |
| ARG-124 | GLN-77 | 2.055 | 29.745 | ion-dipole |
| GLN-143 | ARG-120 | 3.546 | -0.811 | ion-dipole |
| ASP-144 | SER-54 | 3.836 | -2.05 | ion-dipole |
| ALA-91 | VAL-24 | 2.517 | -6.245 | hydrophobic |
| ALA-91 | LEU-27 | 3.167 | -2.694 | hydrophobic |
| ALA-91 | LEU-28 | 3.696 | -5.921 | hydrophobic |
| VAL-94 | VAL-24 | 3.71 | -0.569 | hydrophobic |
| VAL-94 | LEU-28 | 2.481 | -7.335 | hydrophobic |
| VAL-94 | LEU-138 | 2.44 | -9.466 | hydrophobic |
| VAL-94 | ILE-139 | 2.531 | -5.961 | hydrophobic |
| ALA-97 | LEU-138 | 3.034 | -1.285 | hydrophobic |
| MET-98 | LEU-131 | 2.412 | -3.99 | hydrophobic |
| MET-98 | VAL-134 | 2.62 | -7.475 | hydrophobic |
| MET-98 | LEU-138 | 2.823 | -6.057 | hydrophobic |
| PHE-99 | LEU-28 | 3.407 | -2.008 | hydrophobic |
| PHE-99 | LEU-31 | 3.512 | -14.508 | hydrophobic |
| PHE-99 | VAL-32 | 2.442 | -6.712 | hydrophobic |
| PHE-99 | VAL-35 | 1.887 | 5.61 | hydrophobic |
| PHE-114 | LEU-149 | 2.374 | -7.876 | hydrophobic |
| PHE-114 | VAL-189 | 3.851 | -4.061 | hydrophobic |
| VAL-122 | LEU-138 | 3.493 | -6.616 | hydrophobic |
| VAL-122 | ALA-141 | 2.987 | -5.236 | hydrophobic |
| LEU-125 | PHE-71 | 3.157 | -10.117 | hydrophobic |
| LEU-125 | ILE-78 | 2.384 | -7.132 | hydrophobic |
| LEU-125 | LEU-137 | 2.95 | -4.205 | hydrophobic |
| LEU-125 | LEU-138 | 2.344 | -8.393 | hydrophobic |
| LEU-125 | ALA-141 | 2.802 | -1.662 | hydrophobic |
| MET-131 | ALA-67 | 3.721 | -2.215 | hydrophobic |
| LEU-132 | VAL-64 | 3.304 | -6.794 | hydrophobic |
| LEU-132 | ALA-67 | 2.666 | -8.556 | hydrophobic |
| LEU-132 | PHE-71 | 3.355 | -5.234 | hydrophobic |
| LEU-132 | LEU-130 | 2.733 | -6.002 | hydrophobic |
| LEU-132 | LEU-131 | 2.975 | -4.482 | hydrophobic |
| LEU-132 | VAL-134 | 2.177 | -0.358 | hydrophobic |
| ALA-135 | VAL-64 | 2.609 | -6.751 | hydrophobic |
| ALA-135 | ALA-67 | 3.228 | 1.62 | hydrophobic |
| VAL-136 | VAL-64 | 3.051 | -7.922 | hydrophobic |
| VAL-136 | MET-103 | 2.405 | -4.233 | hydrophobic |
| VAL-136 | ALA-127 | 2.416 | -3.731 | hydrophobic |
| VAL-136 | LEU-130 | 3.097 | -0.707 | hydrophobic |
| VAL-136 | LEU-131 | 2.831 | -5.006 | hydrophobic |
| VAL-137 | VAL-35 | 2.518 | -7.077 | hydrophobic |
| LEU-139 | LEU-60 | 2.473 | -14.126 | hydrophobic |
| LEU-139 | ALA-61 | 3.322 | -6.081 | hydrophobic |
| LEU-139 | VAL-64 | 2.469 | -3.839 | hydrophobic |
| LEU-139 | MET-103 | 2.373 | -2.262 | hydrophobic |
| LEU-139 | PHE-110 | 2.71 | -5.46 | hydrophobic |
| LEU-139 | MET-123 | 2.614 | -12.503 | hydrophobic |
| ILE-140 | LEU-38 | 2.985 | -3.762 | hydrophobic |
| ILE-140 | MET-103 | 3.958 | -1.131 | hydrophobic |
| ILE-140 | MET-123 | 3.089 | -5.379 | hydrophobic |
| ILE-140 | VAL-124 | 2.228 | -7.829 | hydrophobic |
| ILE-140 | ALA-127 | 2.431 | -0.312 | hydrophobic |
| ASN-121 | SER-82 | 3.716 | -1.472 | H-bond |
| PHE-114 | ARG-148 | 2.354 | -15.36 | cation-$\pi$ |
| TYR-142 | LYS-56 | 2.007 | 2.157 | cation-$\pi$ |
| TYR-142 | ARG-120 | 3.52 | -7.796 | cation-$\pi$ |

Table S2: **Residue-level interaction analysis of independently docked $\beta$-catenin-IQGAP1 complex**

| $\beta$-catenin Residue | IQGAP1 Residue | Distances (Å) | Energies (KJ/mol) | Interaction type |
|---|---|---|---|---|
| ARG-90 | ASP-70 | 1.915 | -178.181 | Salt bridge |
| ARG-93 | ASP-70 | 1.815 | -213.018 | Salt bridge |
| HIS-118 | GLU-46 | 2.783 | -32.558 | Salt bridge |
| ARG-124 | GLU-45 | 1.816 | -93.396 | Salt bridge |
| ARG-124 | GLU-46 | 1.765 | -106.567 | Salt bridge |
| TYR-86 | GLN-69 | 3.209 | -5.436 | $\pi$-H-bond |
| TYR-86 | GLN-73 | 2.501 | -11.011 | $\pi$-H-bond |
| THR-112 | HIS-67 | 2.348 | -8.141 | $\pi$-H-bond |
| THR-120 | HIS-67 | 3.247 | 0.03 | $\pi$-H-bond |
| GLN-85 | GLU-77 | 2.657 | -51.248 | ion-dipole |
| THR-112 | ASP-70 | 3.564 | -2.032 | ion-dipole |
| THR-120 | ASP-49 | 2.867 | -67.172 | ion-dipole |
| ASN-121 | GLU-46 | 2.239 | -5.502 | ion-dipole |
| ASN-121 | ASP-49 | 1.978 | -95.367 | ion-dipole |
| SER-129 | GLU-62 | 2.401 | -24.005 | ion-dipole |
| LYS-133 | GLN-60 | 2.607 | -6.954 | ion-dipole |
| ALA-87 | LEU-74 | 2.823 | -4.694 | hydrophobic |
| ALA-87 | MET-82 | 2.487 | -0.006 | hydrophobic |
| VAL-94 | PHE-64 | 2.926 | -4.255 | hydrophobic |
| MET-98 | MET-61 | 3.65 | -13.81 | hydrophobic |
| PHE-114 | VAL-47 | 2.74 | -5.627 | hydrophobic |
| VAL-122 | MET-65 | 2.848 | -5.586 | hydrophobic |
| LEU-125 | MET-65 | 2.428 | -5.375 | hydrophobic |
| VAL-136 | VAL-59 | 2.681 | -2.703 | hydrophobic |
| GLN-85 | GLN-73 | 2.881 | -6.498 | H-bond |
| THR-112 | GLN-69 | 2.532 | -2.889 | H-bond |
| ASN-121 | THR-44 | 2.368 | 2.007 | H-bond |
| ARG-90 | PHE-64 | 2.002 | 1.764 | cation-$\pi$ |

Table S3: **IQGAP1-$\beta$-catenin salt bridges formed in the best scoring model of replicates 1, 2, 3 and 4 from the partially blind protocol at pull velocity 12 nm $ns^{-1}$ estimated with a 5 Å cutoff. Here all the residues on $\beta$-catenin are available for docking of IQGAP1 with no specific active residue guidance included.**

| Conformer | Replicate 1 | |
|---|---|---|
| 10 | ARG-124 | GLU-19 |
| | GLU-101 | LYS-11 |
| | GLU-101 | LYS-55 |
| | GLU-127 | LYS-101 |
| | GLU-127 | LYS-20 |
| | HIS-118 | GLU-19 |
| 150 | ASP-115 | LYS-35 |
| | GLU-101 | LYS-102 |
| | GLU-127 | LYS-101 |
| | GLU-127 | LYS-20 |
| | HIS-118 | GLU-19 |
| 200 | ASP-115 | LYS-35 |
| | GLU-101 | LYS-102 |
| | GLU-101 | LYS-11 |
| | GLU-127 | LYS-101 |
| | GLU-127 | LYS-20 |
| | HIS-118 | GLU-19 |
| 250 | ARG-124 | GLU-46 |
| | ARG-93 | GLU-45 |
| | ARG-95 | GLU-51 |
| | GLU-101 | LYS-53 |
| | GLU-127 | HIS-67 |
| | HIS-134 | ASP-49 |
| | HIS-134 | ASP-70 |
| | LYS-133 | GLU-51 |
| | LYS-133 | GLU-62 |
| 400 | GLU-101 | ARG-16 |
| | GLU-101 | LYS-20 |
| | GLU-127 | LYS-101 |
| | HIS-134 | GLU-19 |
| | LYS-133 | GLU-19 |
| 500 | ARG-124 | GLU-62 |
| | ARG-90 | ASP-49 |
| | ARG-90 | GLU-45 |
| | ARG-90 | GLU-46 |
| | ARG-90 | GLU-51 |
| | ARG-95 | ASP-70 |
| | HIS-134 | GLU-77 |
| | LYS-133 | GLU-77 |

| Conformer | Replicate 2 | |
|---|---|---|
| 10 | ASP-115 | HIS-18 |
| | GLU-101 | LYS-11 |
| | GLU-127 | LYS-101 |
| | GLU-127 | LYS-102 |
| | GLU-127 | LYS-20 |
| | HIS-118 | GLU-19 |
| 150 | ARG-124 | ASP-49 |
| | ARG-124 | GLU-46 |
| | ARG-93 | GLU-62 |
| | ASP-115 | LYS-53 |
| | GLU-101 | LYS-83 |
| | GLU-127 | HIS-67 |
| | HIS-118 | GLU-45 |
| | HIS-118 | GLU-46 |
| | HIS-118 | GLU-51 |
| | LYS-133 | GLU-77 |
| 200 | ASP-115 | ARG-16 |
| | ASP-115 | LYS-35 |
| | GLU-101 | LYS-106 |
| | GLU-127 | LYS-101 |
| | GLU-127 | LYS-20 |
| | HIS-118 | GLU-19 |
| 250 | ASP-115 | LYS-35 |
| | GLU-127 | LYS-101 |
| | GLU-127 | LYS-20 |
| | HIS-118 | GLU-19 |
| 400 | ARG-124 | ASP-70 |
| | ARG-124 | GLU-77 |
| | ARG-93 | ASP-86 |
| | ARG-95 | GLU-62 |
| | ASP-83 | ARG-87 |
| | HIS-118 | GLU-77 |
| 500 | ARG-90 | GLU-45 |
| | ARG-90 | GLU-46 |
| | ASP-83 | LYS-106 |
| | GLU-127 | LYS-101 |
| | LYS-133 | GLU-19 |

| Conformer | Replciate 3 | |
|---|---|---|
| 10 | GLU-101 | LYS-11 |
| | GLU-101 | LYS-55 |
| | GLU-127 | LYS-101 |
| | GLU-127 | LYS-20 |
| | HIS-118 | GLU-19 |
| 150 | ASP-115 | LYS-35 |
| | GLU-101 | LYS-102 |
| | GLU-101 | LYS-11 |
| | GLU-127 | LYS-101 |
| | GLU-127 | LYS-20 |
| | HIS-118 | GLU-19 |
| 200 | GLU-101 | LYS-102 |
| | GLU-127 | ARG-16 |
| | GLU-127 | HIS-18 |
| | GLU-127 | LYS-20 |
| 250 | GLU-101 | LYS-11 |
| | GLU-101 | LYS-55 |
| | GLU-127 | LYS-101 |
| | GLU-127 | LYS-102 |
| | GLU-127 | LYS-20 |
| | HIS-118 | GLU-19 |
| 400 | ARG-124 | GLU-62 |
| | ARG-90 | GLU-40 |
| | ARG-90 | GLU-51 |
| | ARG-93 | GLU-51 |
| | ARG-95 | GLU-45 |
| | ASP-83 | LYS-11 |
| | ASP-83 | LYS-53 |
| | ASP-83 | LYS-55 |
| | HIS-118 | GLU-62 |
| 500 | ARG-90 | GLU-40 |
| | ARG-90 | GLU-51 |
| | ARG-95 | ASP-49 |
| | ARG-95 | GLU-46 |
| | GLU-101 | HIS-67 |
| | LYS-133 | ASP-70 |

| Conformer | Replciate 4 | |
|---|---|---|
| 10 | ARG-124 | GLU-46 |
| | ARG-93 | GLU-45 |
| | GLU-101 | LYS-53 |
| | GLU-127 | HIS-67 |
| | HIS-134 | ASP-70 |
| | LYS-133 | GLU-51 |
| 150 | ASP-115 | LYS-35 |
| | GLU-101 | LYS-11 |
| | GLU-127 | LYS-101 |
| | GLU-127 | LYS-20 |
| | HIS-118 | GLU-19 |
| 200 | GLU-101 | LYS-53 |
| | GLU-127 | ARG-16 |
| | GLU-127 | LYS-102 |
| | GLU-127 | LYS-11 |
| | GLU-127 | LYS-20 |
| 250 | ASP-115 | LYS-20 |
| | HIS-118 | GLU-19 |
| 400 | ARG-124 | ASP-49 |
| | ARG-124 | ASP-70 |
| | ARG-95 | ASP-86 |
| 500 | ARG-124 | ASP-70 |
| | ARG-124 | GLU-77 |
| | ARG-90 | ASP-28 |
| | ARG-93 | ASP-86 |
| | ARG-95 | GLU-62 |
| | ASP-83 | ARG-87 |
| | HIS-118 | GLU-77 |

Table S4: **Interfacial residues for the docked IQGAP1 and the conformer 150 of replicate 1 chosen from SMD at pull velocity 12 nm $ns^{-1}$. This results are from docking with active residue guidance.**

| Conformer | $\beta$-catenin | IQGAP1 | Interaction type |
|---|---|---|---|
| | ARG-124 | SER-110 | Ion-dipole |
| | GLN-113 | ASN-100 | H-bond |
| | GLN-113 | TYR-68 | π-H bond |
| | GLN-123 | ASN-100 | H-bond |
| | GLN-123 | LYS-101 | Ion-dipole |
| | GLN-123 | LYS-102 | Ion-dipole |
| | GLN-123 | PHE-97 | π-H bond |
| | GLN-130 | LYS-102 | Ion-dipole |
| 150 | GLU-101 | ARG-16 | Salt bridge |
| | GLU-101 | LYS-20 | Salt bridge |
| | GLU-127 | LYS-102 | Salt bridge |
| | HIS-118 | SER-110 | Ion-dipole |
| | SER-111 | LYS-101 | Ion-dipole |
| | SER-111 | PHE-97 | π-H bond |
| | SER-129 | LYS-101 | Ion-dipole |
| | THR-102 | GLU-19 | Ion-dipole |
| | THR-102 | LYS-20 | Ion-dipole |

Table S5: **Interfacial residues for the docked IQGAP1 and the conformer 200 of replicate 1 chosen from SMD at pull velocity 12 nm $ns^{-1}$. This results are from docking with active residue guidance.**

| Conformer | $\beta$-catenin | IQGAP1 | Interaction type |
|---|---|---|---|
| 200 | ARG-124 | SER-110 | Ion-dipole |
| | ARG-124 | TYR-104 | Ion-dipole |
| | GLN-113 | ASN-100 | H-bond |
| | GLN-113 | TYR-68 | π-H bond |
| | GLN-123 | ASN-100 | H-bond |
| | GLN-123 | LYS-101 | Ion-dipole |
| | GLN-123 | PHE-97 | π-H bond |
| | GLN-130 | LYS-101 | Ion-dipole |
| | GLN-130 | LYS-102 | Ion-dipole |
| | GLU-101 | ARG-16 | Salt bridge |
| | GLU-101 | LYS-20 | Salt bridge |
| | GLU-127 | LYS-101 | Salt bridge |
| | GLU-127 | LYS-102 | Salt bridge |
| | HIS-118 | SER-110 | π-H bond |
| | HIS-118 | TYR-104 | cation-π |
| | SER-111 | LYS-101 | Ion-dipole |
| | SER-111 | PHE-97 | π-H bond |
| | THR-102 | LYS-20 | Ion-dipole |

Table S6: **Interfacial residues for the docked IQGAP1 and the conformer 400 of replicate 1 chosen from SMD at pull velocity 12 nm $ns^{-1}$. This results are from docking with active residue guidance.**

| Conformer | $\beta$-catenin | IQGAP1 | Interaction type |
|---|---|---|---|
| 400 | ARG-90 | GLU-45 | Salt bridge |
| | ARG-90 | GLU-46 | Salt bridge |
| | ARG-90 | PHE-103 | cation-π |
| | ARG-90 | THR-44 | Ion-dipole |
| | ARG-93 | SER-107 | Ion-dipole |
| | ASN-121 | ASN-100 | H-bond |
| | ASN-121 | TYR-104 | π-H bond |
| | ASN-141 | GLU-19 | Ion-dipole |
| | ASP-115 | SER-110 | Ion-dipole |
| | ASP-83 | LYS-106 | Salt bridge |
| | ASP-83 | SER-6 | Ion-dipole |
| | GLN-85 | SER-3 | H-bond |
| | GLN-85 | SER-5 | H-bond |
| | GLU-101 | ARG-16 | Salt bridge |
| | HIS-118 | SER-110 | π-H bond |
| | PHE-114 | SER-110 | π-H bond |
| | PHE-99 | ARG-16 | cation-π |
| | PHE-99 | LYS-20 | cation-π |
| | SER-129 | LYS-101 | Ion-dipole |
| | SER-129 | PHE-97 | π-H bond |
| | THR-112 | LYS-106 | Ion-dipole |
| | THR-112 | SER-107 | H-bond |
| | TYR-86 | LYS-106 | cation-π |
| | TYR-86 | SER-107 | π-H bond |

Table S7: **IQGAP1-$\beta$-catenin salt bridges formed in the best scoring model of replicates 1, 2, 3 and 4 from the restrained protocol at pull velocity 12 nm $ns^{-1}$ estimated with a 5 Å cutoff. Here active residue guidance is included.**

| Conformer | Replicate 1 | | Conformer | Replicate 2 | | Conformer | Replciate 3 | | Conformer | Replciate 4 | |
|---|---|---|---|---|---|---|---|---|---|---|---|
| 10 | GLU-101 | ARG-16 | 10 | GLU-101 | ARG-16 | 10 | GLU-101 | ARG-16 | 10 | GLU-101 | ARG-16 |
| | GLU-101 | LYS-20 | | GLU-101 | LYS-20 | | GLU-101 | LYS-20 | | GLU-101 | LYS-20 |
| | GLU-127 | LYS-101 | | GLU-127 | LYS-101 | | GLU-127 | LYS-102 | | GLU-127 | LYS-102 |
| | GLU-127 | LYS-102 | | GLU-127 | LYS-102 | 150 | GLU-101 | ARG-16 | 150 | GLU-101 | ARG-16 |
| 150 | GLU-101 | ARG-16 | 150 | GLU-101 | ARG-16 | | GLU-101 | LYS-20 | | GLU-101 | LYS-20 |
| | GLU-101 | LYS-20 | | GLU-101 | LYS-20 | | GLU-127 | LYS-101 | | GLU-127 | LYS-102 |
| | GLU-127 | LYS-102 | | GLU-127 | LYS-101 | | GLU-127 | LYS-102 | 250 | ARG-90 | GLU-45 |
| 250 | ARG-90 | GLU-45 | | GLU-127 | LYS-102 | 250 | ARG-90 | GLU-45 | | ARG-90 | GLU-46 |
| | ARG-90 | GLU-46 | 250 | ARG-90 | GLU-45 | | ARG-90 | GLU-46 | | ASP-83 | LYS-106 |
| | ASP-83 | LYS-106 | | ARG-90 | GLU-46 | | ASP-83 | LYS-106 | | GLU-101 | ARG-16 |
| 400 | ARG-90 | GLU-45 | | ASP-83 | LYS-106 | | GLU-101 | ARG-16 | 400 | ARG-90 | GLU-45 |
| | ARG-90 | GLU-46 | | GLU-101 | ARG-16 | | LYS-133 | GLU-19 | | ARG-90 | GLU-46 |
| | ASP-83 | LYS-106 | | GLU-101 | LYS-20 | 400 | ARG-90 | GLU-45 | | ASP-83 | LYS-106 |
| | GLU-101 | ARG-16 | 400 | ARG-90 | GLU-45 | | ARG-90 | GLU-46 | | GLU-127 | LYS-101 |
| 500 | ARG-90 | GLU-45 | | ARG-90 | GLU-46 | | ASP-83 | LYS-106 | | LYS-133 | GLU-19 |
| | ARG-90 | GLU-46 | | ASP-83 | LYS-106 | | GLU-101 | ARG-16 | 500 | ARG-90 | GLU-45 |
| | ASP-83 | LYS-106 | | GLU-101 | ARG-16 | | LYS-133 | GLU-19 | | ARG-90 | GLU-46 |
| | GLU-101 | ARG-16 | | GLU-127 | LYS-101 | 500 | ARG-90 | GLU-45 | | | |
| | LYS-133 | GLU-19 | | LYS-133 | GLU-19 | | ARG-90 | GLU-46 | | | |
| | | | 500 | ARG-90 | GLU-45 | | ASP-83 | LYS-106 | | | |
| | | | | ARG-90 | GLU-46 | | GLU-101 | ARG-16 | | | |
| | | | | ASP-83 | LYS-106 | | | | | | |
| | | | | GLU-101 | ARG-16 | | | | | | |
| | | | | GLU-127 | LYS-101 | | | | | | |
| | | | | LYS-133 | GLU-19 | | | | | | |

Vymětal, Jiří, David Jakubec, Jakub Galgonek, and Jiří Vondrášek. 2021. "Amino Acid Interactions (INTAA) web server v2.0: a single service for computation of energetics and conservation in biomolecular 3D structures." *Nucleic acids research* 49 (W1): W15–W20.
